# Subcellular transcriptome of radial glia reveals compartmentalized control of cortical development

**DOI:** 10.1101/2025.10.10.679263

**Authors:** Brooke R. D’Arcy, Camila Manso Musso, Chia-Fang Lee, Lucas D. Serdar, Stephany Perez-Sanchez, Virginia Fernández, Vίctor Borrell, Debra L. Silver

**Author notes:** Authors contributed equally to the work (Brooke R. D’Arcy, Camila Manso Musso).

## Abstract

RNA localization and local translation mediate spatial and temporal control of polarized cells, including radial glial cells (RGCs) which produce and organize neurons and glia. Within RGCs, RNAs are transported long distances to basal endfeet, where they can undergo local translation. However, the subcellular composition of RGCs and function of local gene regulation remains largely unknown. Here, we discover that basal endfeet harbor a rich transcriptome including a Dynein component critical for subcellular RGC function. By purifying RGC compartments *in vivo*, we discover ∼3000 endfoot transcripts, including ∼800 highly enriched compared to cell bodies. Many endfoot-enriched transcripts exhibit conserved subcellular localization in neurons and glia and are associated with neurodevelopmental disease. We show that endfoot-enriched *Dync1li2* regulates RGC basal morphology and subsequently interneuron organization. Finally, we develop LOCAL-KD, a CRISPR-Cas13 based method for subcellular mRNA knockdown *in vivo*. Leveraging this, we demonstrate that endfoot-localized *Dync1li2* is critical for RGC morphology. Our study establishes experimental paradigms to understand RNA localization in the nervous system. Moreover, we discover RGCs have a vast subcellular transcriptome, revealing foundational insights into how RGCs control cortical development.

## Introduction

Local mRNA regulation is an essential layer of gene expression in polarized cells. mRNA localization coupled with local translation provides precise spatial and temporal control of proteomes within subcellular compartments. In neurons, localized mRNA transcripts and translation underlie many fundamental cellular processes, such as axon guidance, dendrite and arbor branching, repair, and synaptic activity [1]. In oligodendrocytes and astrocytes, local translation is implicated in diverse processes including morphology and myelination [2]. An emerging model for RNA localization in the developing cortex are highly polarized neural progenitors termed radial glial cells (RGCs).

RGCs are responsible for generating neurons and glial cells, and for guiding them to their proper destination [3–5]. They are bipolar, with cell bodies positioned adjacent to the ventricle, and a long basal process that attaches to the pial surface by basal endfeet (**Fig. 1A**). These basal structures are also found in outer radial glia (oRGC/bRGC), which are prominent in gyrencephalic species[6–8]. The basal process can extend hundreds of micrometers in mice and centimeters in humans and serves as a scaffold for migration of newborn excitatory neurons into the cortical plate [9, 10]. Basal endfeet are tightly connected to a basement membrane (BM) forming a physical barrier between the brain and overlying meninges [11]. Importantly, basal endfeet are embedded in a unique niche composed of overlying meninges, as well as interneurons and Cajal-Retzius cells of the marginal zone (MZ) (**Fig. 1A**). As development proceeds, RGC basal branches become more morphologically complex and endfoot number increases [12]. Moreover, disruptions to RGC morphology and basal lamina integrity can lead to neurodevelopmental diseases, such as cobblestone malformation and lissencephaly [13, 14]. While this reinforces the central role of RGC morphology in cortical development, how this subcellular architecture is molecularly controlled remains largely elusive.

**Fig. 1:**
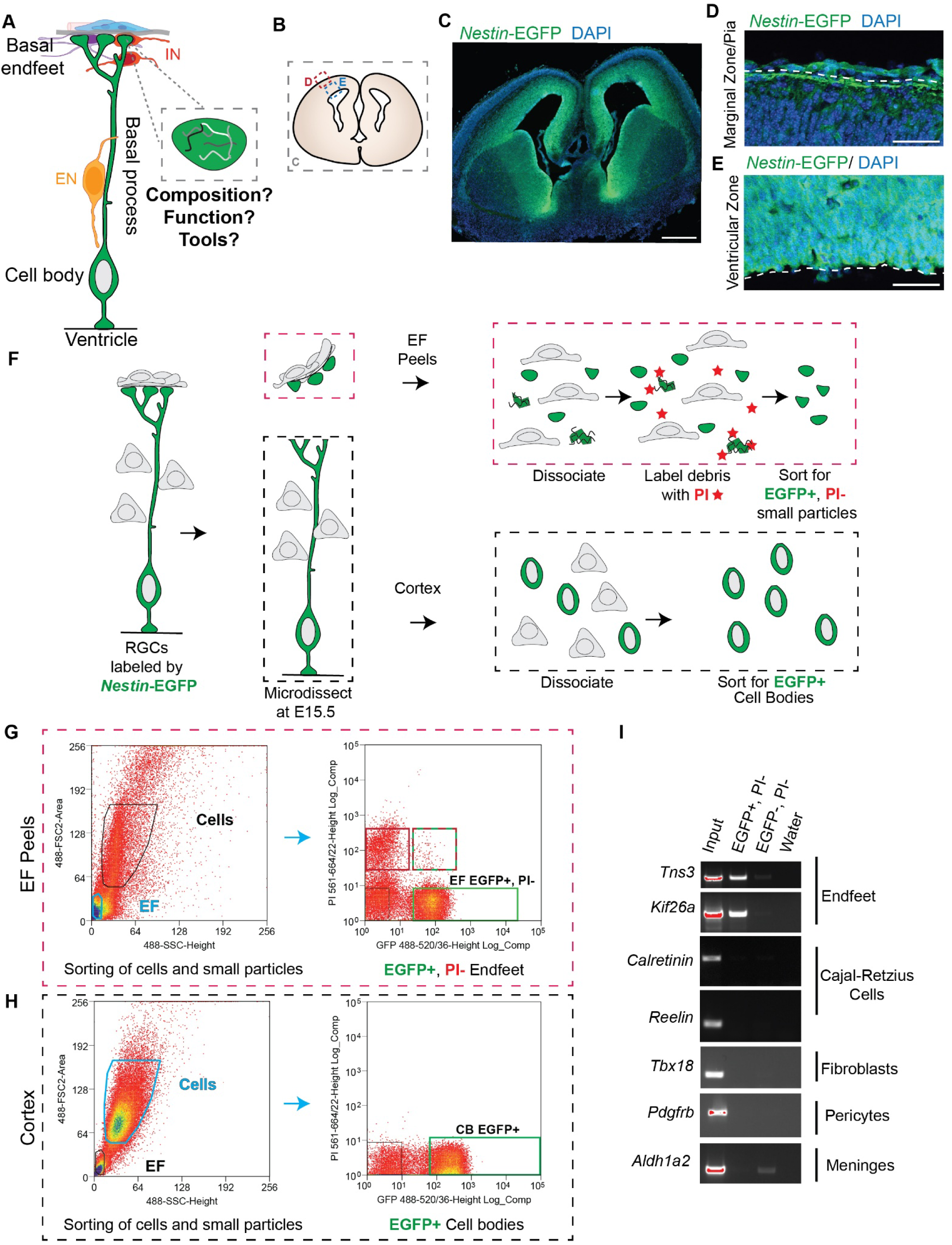
A method for isolation of RGC basal endfeet from embryonic cortices. **(A)** Cartoon representation of RGC morphology, RGC niche and key questions of the study. Fibroblasts (blue), blood vessels (pink), basement membrane (grey), Cajal-Retzius cell (purple), interneuron (red, IN), excitatory neuron (yellow, EN). **(B)** Cartoon representation of a coronal brain section and position of images in C-E. **(C)** Expression of *Nestin*-EGFP (green) in coronal sections of E15.5 brains co-stained with DAPI (blue). **(D)** *Nestin*-EGFP (green) and DAPI (blue) stained sections of the marginal zone/pia with EGFP+ endfeet. White dotted line denotes the endfoot/pia boundary. **(E)** *Nestin*-EGFP (green) and DAPI (blue) stained sections of the ventricular zone with EGFP+RGC cell bodies. **(F)** Cartoon representation of the microdissection and FACS method to isolate RGC cell bodies (bottom) and basal endfeet (EF, top) from embryonic mouse brains. Samples collected at E15.5. **(G)** Example FACS plots for sorting basal endfeet. Separation of cells and small particles by size (left) and separation of small particles into debris and endfeet by fluorescence (right). Green box represents EGFP+, propidium iodide (PI) - Endfeet collected for RNA isolation. **(H)** Example FACS plots for sorting cell bodies. Separation of cells and small particles by size (left) and separation of cells into EGFP+ and EGFP-by fluorescence (right). Green box represents EGFP+ cell bodies collected for RNA isolation. **(I)** RT-PCR for marker genes (left) of cell types and compartments (right) in the input, EGFP+ and EGFP-samples for endfoot peel samples. Oversaturated pixels pseudo colored red. Scale bars: 500µm (C), 50µm (D and E).

Within RGC basal processes, mRNA is trafficked to basal endfeet, where mRNAs can localize and undergo local translation [15]. Asymmetric distribution of mRNAs in RGCs has been hypothesized to influence critical features of cortical development, such as basement membrane integrity and cell fate [14, 16, 17]. Further, localized transcripts are required for local RGC morphology and attachment to the BM. For example, while depletion of *Arhgap11a* or non-muscle myosin heavy chain *Myh9* reduces RGC branching complexity, loss of *Myh10* impairs endfoot adhesion [18, 19]. These disruptions to RGC morphology have non-cell autonomous impacts on interneuron organization and number. Towards understanding the subcellular composition of RGCs, BioID experiments *in vivo* have uncovered a proteome of RGC basal endfeet including proteins related to ECM, actin and microtubule cytoskeletal regulation, and ubiquitination [18]. Further, FMRP RNA immunoprecipitations have shown over 100 transcripts in endfeet [15]. However, a comprehensive atlas of the RGC subcellular transcriptome is lacking which impedes our ability to understand the function of RNA localization in RGCs [15–21].

In this study, we develop a method for purifying RGC cell bodies and basal endfeet from the developing mouse brain, which we leverage to define subcellular transcriptomes of endfeet. We discover over 3,000 endfoot localized genes, 800 of which are highly enriched in endfeet compared to cell bodies, including many with conserved subcellular localization in other cell types and linked to disease. We functionally interrogate roles for dynein light intermediate chain 2 (*Dync1li2*), which is among the most enriched transcripts in endfeet. We demonstrate that *Dync1li2* is essential for regulating RGC morphology and interneuron organization during cortical development. Furthermore, we develop a CRISPR-Cas13d based method for spatial manipulation of mRNAs in RGCs *in vivo*, which we use to demonstrate subcellular requirements of *Dync1li2* for RGC basal morphology. Our study establishes experimental paradigms to understand mRNA localization, uncovering new principles of gene regulation in RGCs. Moreover, this transcriptome generates new hypotheses about endfoot function in the developing cortex.

## Results

### Development of a purification method for RGC basal endfeet and cell bodies *in vivo*

To expand our understanding of RGC functionality and local gene regulation, we aimed to identify a global, unbiased transcriptome of RGC basal endfeet *in vivo*. Towards this, we needed to develop a new approach to isolate and purify this subcellular compartment from the rest of the embryonic brain. First, we devised a method to distinguish RGCs and their endfeet from other cells in the cortex. To achieve this, we fluorescently labeled RGCs *in vivo* using *Nestin*-EGFP transgenic mice [22]. In these mice, EGFP expression is driven by the *Nestin* promoter and second intron, which acts as an enhancer in neural cells [22]. *Nestin* encodes an intermediate filament protein which robustly marks RGCs. Consistent with this, we observed labeling of RGC cell bodies, basal processes, and endfeet with EGFP at E15.5 (**Fig. 1B-1E**). This approach enabled us to efficiently label all the RGCs in the developing cortex.

We next developed methods to separate the EGFP+ basal endfeet from their respective cell bodies. To achieve this, we first peeled the meninges from the embryonic cortex as previously termed endfoot peels [15], and then separately dissociated the meninges tissue (including endfeet) and underlying cortex (including RGC cell bodies) (**Fig. 1F**). The EGFP labeling from the *Nestin*-EGFP mice made it possible to then isolate EGFP+ cells and endfeet from their corresponding tissues by fluorescence activated cell sorting (FACS). As endfeet are approximately the same size as debris from tissue dissociation, we used propidium iodide (PI) to sort out debris as it labels exposed nucleic acids but not intact cell membranes (**Fig. 1F**). The cells were sorted using standard gating for both size and EGFP while the endfoot samples were sorted for EGFP positive, PI negative small particles (5-10µm) (**Fig. 1G-1H**).

We next assessed the purity of sorted endfeet by measuring expression of known endfoot localized transcripts as well as transcripts from surrounding cortical and meninges cells. In sorted endfeet, using RT-PCR, we detected mRNAs for *Tns3* and *Kif26a*, RGC markers which had previously been shown to localize to endfeet (**Fig. 1I**) [15]. Markers for Cajal-Retzius cells, fibroblasts, pericytes, and meninges were detected only in endfeet peel input, but not in purified endfeet (**Fig. 1I**). RT-PCR of these same markers revealed that propidium iodide treatment is necessary for generating pure endfeet (**Fig. S1A).** From this we conclude that sorting *Nestin*-EGFP positive, PI negative small particles, isolates RGC basal endfeet with minimal contamination from surrounding cells of the MZ and pia.

### Discovery of a global, subcellular transcriptome of RGC basal endfeet

Using the new method described above, we collected RGC basal endfeet and their corresponding cell bodies for RNA sequencing. E15.5 was chosen as it is a mid-stage of neurogenesis and complements previous analyses of local proteomes and local translation at the same stage [15, 18]. Three biological replicates were performed with multiple embryos (from 3-4 litters) pooled per replicate. In total, over 300,000 endfeet were collected for each biological replicate. The endfoot and cell body replicates clustered separately by PCA (**Fig. S2B**). Overall, our experiment identified 25,484 unique transcripts across all samples. 7,726 of these transcripts had significant differential localization in endfeet and cell bodies (**Fig. 2A-2B, Table S1**).

**Fig. 2:**
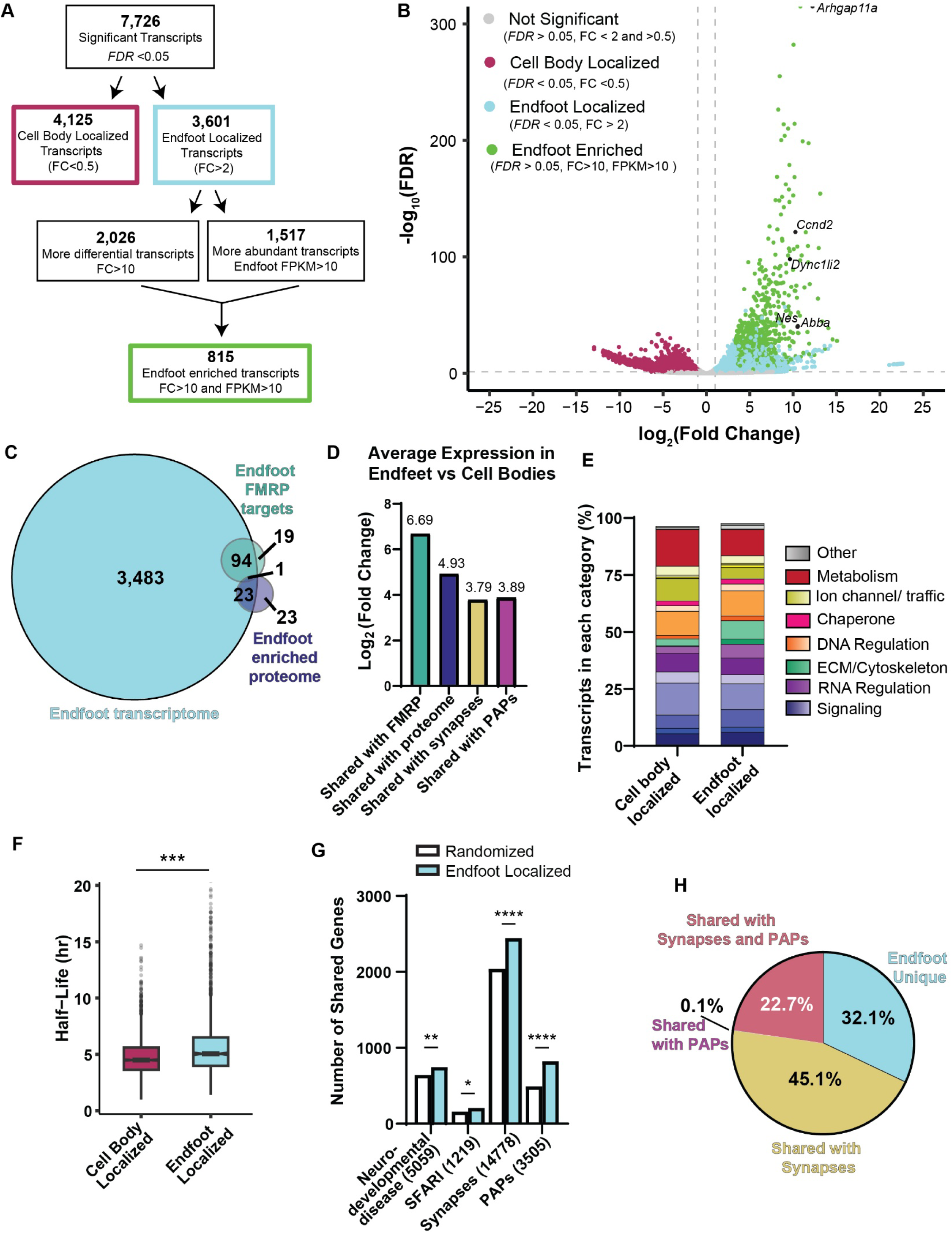
Global transcriptome of endfeet reveals diverse, subcellular mRNA composition. **(A)** Criteria used to identify statistically significant transcripts, and those categorized as cell body localized (maroon), endfoot localized (light blue), or endfoot enriched (green) along with the number of genes identified in each category. n= 3 biological replicates for endfeet and cell bodies. 10-20 brains were pooled per replicate totaling 331,000-355,000 endfeet and 150,000-200,000 cell bodies from 13 litters. **(B)** Volcano plot comparing – log_10_(FDR value) and log_2_(FC) for all transcripts identified in endfoot and cell body samples by RNA sequencing. Points represent transcripts that are not significant (grey), significantly localized to cell bodies (maroon), significantly localized to endfeet (light blue), or endfoot enriched (green). Horizontal grey dotted line denotes filter for statistical significance (FDR<0.05). Vertical grey dotted lines denote filters for FC significance (FC> 2 for endfeet and FC< 2 for cell bodies). **(C)** Venn diagram comparing transcripts identified as endfoot localized (light blue) to previously published endfoot data sets, showing overlap with FMRP targets in endfeet (teal) and endfoot enriched proteins (indigo). **(D)** Average Log_2_(FC) for all transcripts in each category in endfeet versus cell bodies. **(E)** Stacked bar graph representing the % of transcripts in each functional category for cell body and endfoot localized transcripts. **(F)** Comparison of mRNA half-lives between cell body (maroon) and endfoot (light blue) localized transcripts. n=2713 cell body transcripts and 1630 endfoot transcripts. **(G)** Bootstrap comparison of the number of endfoot localized (light blue) or randomized (white) transcripts shared with each dataset. Compared to neurodevelopmental disease genes, SFARI genes, synapse localized transcripts and peripheral astrocyte process (PAP) localized transcripts. **(H)** Pie chart comparing the fraction of endfoot localized transcripts that are uniquely localized to endfeet (light blue), localized to synapses (yellow), localized to PAPs (magenta), and localized to both synapses and PAPs (salmon). (A, B, E) Stats: (F) Student’s unpaired, two-tailed t test, (G) two-tailed Fisher’s exact test. ns p-value > 0.05, *p-value < 0.05, **p-value < 0.01, ***p-value < 0.001, ****p-value < 0.0001.

We next probed this dataset for purity by assessing the expression of known markers of endfeet and potentially contaminating cell types. Amongst the most enriched and abundant transcripts within basal endfeet were *Ccnd2*, *Abba*, *Nestin*, and *Arhgap11a,* which showed 1,769, 2,173, 2,051, and 4,707-fold enrichment compared to the cell body, respectively (**Fig. 2B**). These transcripts were previously shown to localize and function in endfeet, bolstering confidence in our dataset [16–19, 21]. We also assessed the purity of sorted endfeet by examining expression of known markers of cells surrounding RGC basal endfeet. For this we generated lists of identifying transcripts for Cajal-Retzius cells, meninges, and cortical interneurons, as defined by previously published datasets [23–25]. All identifying transcripts had low FPKM values or were undetected in endfeet (**Fig. S1C-E**). Additionally, their fold changes were close to zero, indicating they are de-enriched in endfeet compared to the cell type of interest (**Fig. S1C-E**). These data reinforce RT-PCR analyses and indicate that purified RGC basal endfeet do not have significant contamination from surrounding cells.

Given these analyses, we classified transcripts according to their subcellular abundance and fold enrichment. We used rigorous cutoffs in order to define transcripts that were localized in either endfoot or cell body (termed endfoot or cell body localized) and those which were highly enriched and abundant in endfeet (termed endfoot enriched). 7,726 transcripts were differentially localized in endfeet versus cell bodies with a fold change (FC) of 2 and FDR<0.05 (**Fig. 2A-2B**, Table S1). Amongst these, 4,125 transcripts were cell body localized (maroon) with FC<2 compared to the endfeet. 3,601 were endfoot localized (light blue) with a FC>2 compared to the cell bodies (**Fig. 2A-2B**). Of the 3,601 endfoot localized transcripts, 2,026 had a FC>10 compared to cell bodies indicating they are more differentially localized in this compartment. 1,517 had a FPKM >10, indicating they are abundant in endfeet (**Fig. 2A**). 815 transcripts were classified as endfoot enriched, with FPKM>10 and FC>10 (**Fig. 2A-2B, Table S1**). In subsequent analyses we refer to the 3,601 (FC>2) as endfoot-localized and the 815 (FC>10 and FPKM>10) as endfoot-enriched.

We observed a wide range of abundances in endfoot localized transcripts, with a maximum FPKM of 51,459.53 for *Ccnd2* and a minimum of 9.56x10^-4^ for *Asxl3*. Further, some transcripts were more prevalent in the cell bodies or in endfeet, by up to 7,968-fold and 17,937-fold, respectively. Therefore, to validate our dataset, we selected transcripts with varying levels of abundance with FPKMs ranging from 4 to 1,054 and enrichment from 11 to 7,131-fold. We validated endfoot expression of *Myh9, Myh10, Myh14,* which were previously shown to localize to endfeet, and *Itsn*, using RT-PCRs of endfoot peels (**Fig. S1F**) [18]. We also validated the localization of 3 endfoot localized transcripts, *Gab2, Mmp14, and Sh3pxd2a,* by smiFISH of *Nestin*-EGFP tissue sections (**Fig. S1G**). Thus, even endfoot transcripts with relatively low mRNA levels and enrichment were confirmed independently.

Following this validation we next focused on analysis of the 3,601 endfoot localized transcripts (**Fig. 2B, light blue)**. We first examined the extent of overlap of endfoot localized transcripts with previously identified transcripts in this compartment. Probing a dataset of 114 FMRP-bound transcripts in endfeet, 95 were found in the endfoot localized transcriptome (**Fig. 2C, Table S5**) [15]. This high degree of overlap validates both datasets. The 95 shared transcripts had an average 6.69 log_2_(FC) in endfeet versus cell bodies (**Fig. 2D**). We next assessed the extent of overlap between the endfoot transcriptome and the previously published endfoot proteome [18]. Of the 47 significantly enriched endfoot proteins, 24 were present at the mRNA level (**Fig. 2C, Table S5**). We observed an average 4.93 log_2_(FC) in endfeet for transcripts shared with this proteome (**Fig. 2D**).Among the shared genes were non-muscle myosins (*Myh9*, *Myh10*, *Myh14*), previously characterized in endfeet [18], as well as kinesins, and collagens. The remaining proteins not found in the transcriptome may be produced outside of endfeet. Altogether, this analysis validates previous investigations of endfoot composition and dramatically expands the dataset.

To understand the types of RNAs found in RGC compartments, we next classified protein ontology. PANTHER revealed the complexity of the endfoot transcriptome with all gene categories reported in cell bodies also found in endfeet (**Fig. 2E, Table S2**). These shared categories were represented in different proportions across compartments (**Fig. 2E, S2A-S2B**). Top protein classes in both populations included metabolite conversion enzymes, gene-specific transcriptional regulators, and protein modifying enzymes. A greater fraction of endfoot transcripts encoded extracellular matrix proteins, cytoskeletal proteins, and defense/immunity proteins (broken down in **Fig. S2B**). The cell body transcripts had a higher percentage of transporters, metabolite interconversion enzymes, and protein modifying enzymes (**Fig. S2B**). These data indicate diverse mRNA composition of RGCs in both cell bodies and endfeet.

We next assessed if the differentially localized transcripts showed distinct RNA stability features. In neurons, neurite localized mRNAs are more stable than those in the cell body [26]. Using SLAM-seq data from the E14.5 cortex [27], we compared the stability of cell body and endfoot localized transcripts. Of the 7,726 differentially localized transcripts, 4,343 had calculated half-lives from this dataset. Amongst these transcripts, 2,713 were cell body localized and 1,630 were endfoot localized (**Fig. S3A-S3C, Table S3**). We then calculated the average half-life for each population based on the values experimentally determined by SLAM-seq [27]. This revealed that endfoot localized transcripts had significantly longer half-lives than their cell body localized counterparts (**Fig. 2F**). This suggests that some endfoot localized mRNAs are more stable than cell body localized mRNAs.

### The endfoot transcriptome significantly overlaps with transcripts associated with neurodevelopmental disease and subcellular compartments of neurons and astrocytes

Mining of databases revealed that many of the 3,601 endfoot localized genes are implicated in neurodevelopmental diseases. Endfoot localized transcripts were represented in datasets of autism, neurodevelopmental disorders, epilepsy, microcephaly, and macrocephaly genes (**Fig. S3D, Table S4**) [28, 29]. Developmental Brain Disorders, SPARK, and Head Circumference showed the most overlap with the endfoot transcriptome, with 32.9, 23.9, and 23.3%, respectively (**Fig. S3D**). Comparison of endfoot localized transcripts with the combined list of disease genes (**Fig. S3D, Table S4**) demonstrated significant overlap compared to randomized control (**Fig. 2G**). Significant overlap was also observed with the SFARI database of Autism associated genes (**Fig. 2G**). This suggests the functional importance of endfoot localized transcripts in brain development.

We next assessed overlap of the 3,601 endfoot localized transcripts with subcellular transcriptomes of neuronal synapses and astrocyte projections [30–34] (**Fig. S3E**). Transcripts overlapping with synapses and peripheral astrocyte processes (PAPs) were highly enriched in endfeet versus cell body, with 3.79 and 3.89 log_2_(FC), respectively (**Fig. 2D**). 2,444 transcripts were shared between the endfoot localized transcriptome and at least one of two neuronal synapse transcriptomes (2,168 shared between all) (**Fig. S3F, Table S6**) [30, 32]. We also compared endfoot localized transcripts to transcripts and proteins localized to astrocyte processes (**Fig. S3G**). Of the 3,505 localized genes in astrocyte processes, 825 were also present in endfeet, representing a 24% overlap (**Fig. S3G, Table S6**) [32–36]. In total, 32.1% of endfoot localized transcripts are uniquely localized in RGCs. In comparison, 45.1% of endfoot localized transcripts are shared with synapses and endfeet, 22.7% are in common with synapses and PAPs, and 0.1% are shared with only PAPs (**Fig. 2H**). The observed overlap was significant between endfoot localized transcripts, synapse localized transcripts, and PAP localized transcripts, compared to a randomized control dataset of equal size (**Fig. 2G**). These data demonstrate that transcripts within endfeet are also localized within protrusions in neurons and astrocytes. Overall, this supports the hypothesis that there are cohorts of transcripts conserved between subcellular compartments of cells of the nervous system [20, 37, 38]. Furthermore, there are transcripts uniquely localized to endfeet suggesting these may have RGC specific subcellular functions. This reinforces the value of using RGCs as a model for understanding mRNA localization.

### Over 800 transcripts are highly enriched in RGC basal endfeet compared to cell bodies

We next focused on transcripts which were the most significantly enriched and abundant in endfeet compared to cell bodies (FC>10 and FPKM>10) (**Fig. 2A**, green). First, we explored if the 815 endfoot enriched transcripts exhibited cis features associated with localization and mRNA metabolism in other cell types, including transcript length and GC content of the endfoot enriched transcripts [39, 40]. Comparisons were made between cell body enriched (control) transcripts (FPKM in cell body >10 and FC <0.5) and endfoot enriched transcripts (FPKM in endfeet >10, FC >10) for their 5′ UTR, CDS, 3′ UTR, and entire mRNA, using FeatureReachR [41]. Across all of these elements, there was a slight but significant 1-2% decrease in GC content in endfoot enriched transcripts compared to control (**Fig. S4C-S4F**). The CDS and 3′ UTRs of endfoot enriched transcripts were significantly longer than their control (cell body enriched) counterparts (**Fig. 3A-3B**). The mean CDS length for endfoot enriched transcripts was 1,744 nucleotides (nts) compared to 1,129 nts for control transcripts, reflecting a 54.6% increase in CDS length for endfoot enriched transcripts (**Fig. 3A**). Similarly, the mean 3′ UTR length for endfoot enriched and control transcripts was 1,916 nts and 1,365 nts, respectively, indicating a 40% increase in endfeet (**Fig. 3B**). Consistent with this, there was an overall increase in total transcript length in endfeet (**Fig. S4A**). In contrast, there was not a significant difference for 5′ UTR length between the two groups (**Fig. S4B**). Longer 3′ UTRs may contain more cis elements to promote mRNA localization and increased CDS length may reflect the energetic benefit of transporting mRNAs for local translation versus trafficking large proteins over long distances.

**Fig. 3.**
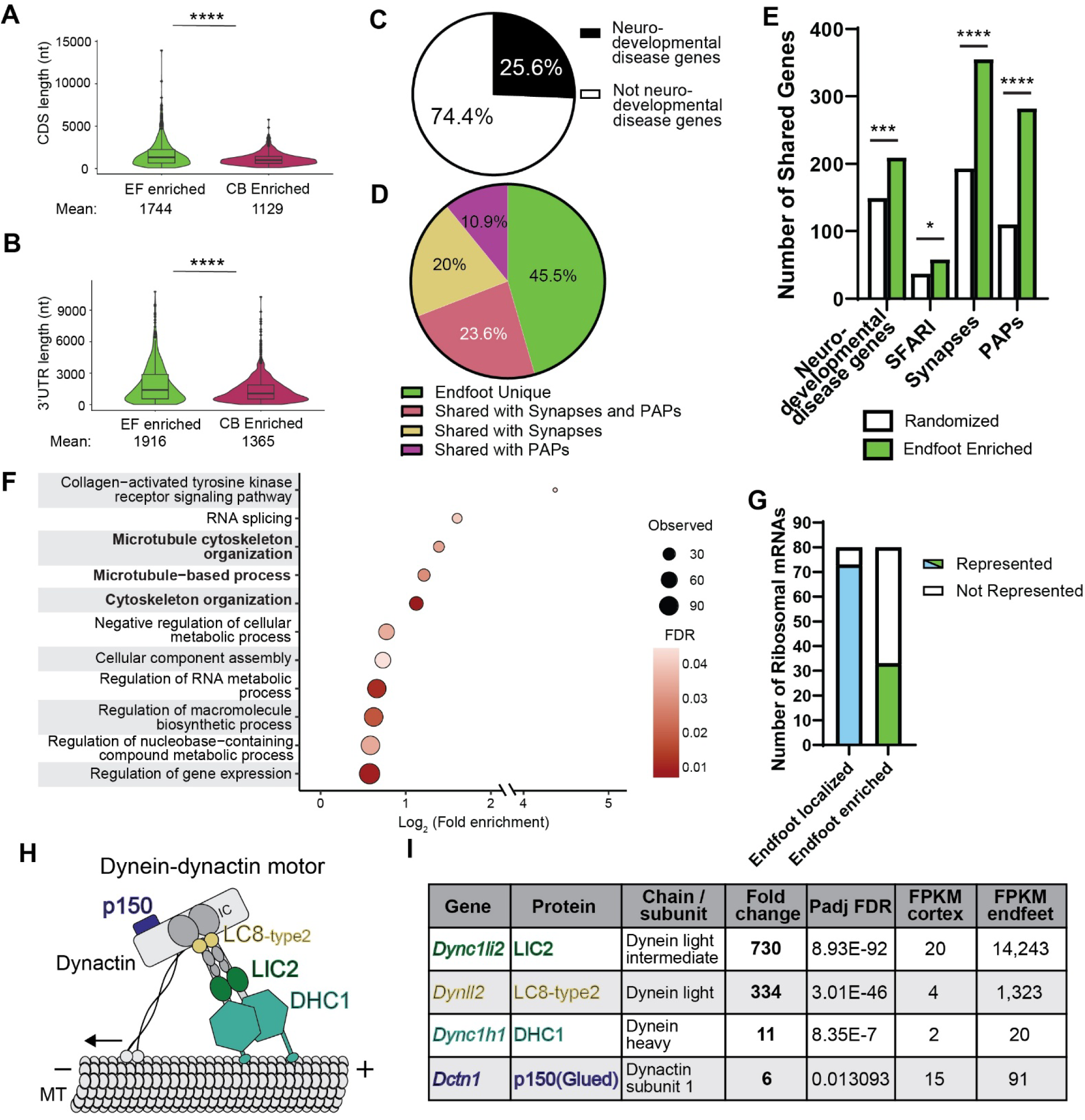
Over 800 endfoot enriched transcripts encode diverse protein classes including dynein components. **(A-B)** Comparison of CDS length **(A)** and 3′ UTR length **(B)** for endfoot enriched genes (EF enriched) (green) and cell body enriched genes (maroon). Cell body enriched genes were classified as those with p-value < 0.05 and FC < -1. Analysis performed using FeatureReachR package in R [41]. **(C)** Pie chart showing the fraction of endfoot enriched transcripts that are represented in neurodevelopmental disease data sets (black) or as yet unknown (white). Disease data sets from Fig. 2E were combined. **(D)** Pie chart comparing the fraction of endfoot enriched transcripts that are uniquely enriched in endfeet (green), enriched in synapses (yellow), enriched in PAPs (magenta), and both synapses and PAPs (salmon). **(E)** Bootstrap comparison of the number of endfoot enriched (green) or randomized (white) transcripts shared with each dataset. Compared to neurodevelopmental disease genes, SFARI genes, synapse localized transcripts and peripheral astrocyte process (PAP) localized transcripts. **(F)** GO Biological Processes analysis of endfoot enriched transcripts. Categories of interest bolded. **(G)** Number of ribosomal RNAs that are endfoot localized (light blue) and endfoot enriched (green). **(H)** Cartoon representation of the dynein-dynactin complex bound to microtubules. **(I)** Table depicting the endfoot enriched transcripts that encode subunits of the dynein-dynactin complex. Stats: (A, B) Wilcoxon rank-sum test. (E) two-tailed Fisher’s exact test. ns p-value > 0.05, *p-value < 0.05, **p-value < 0.01, ***p-value < 0.001, ****p-value < 0.0001. MT, microtubules; HC, heavy chain; IC, intermediate chain, LIC, light intermediate chain, LC, light chain; Robl, Roadblock.

We next assessed the composition of the endfoot enriched transcriptome. One quarter of endfoot enriched transcripts were implicated in neurodevelopmental diseases, supporting their functional roles in brain development (**Fig. 3C, Table S8**). A subset of endfoot enriched transcripts were also localized in subcellular compartments of neurons and astrocytes. Among the 815 endfoot enriched transcripts, 44.5% were uniquely localized in RGCs (**Fig. 3D, Table S9**). 18.2%, 11.9%, and 25.4% of transcripts were also shared with synapses, PAPs, and all 3 compartments respectively. The observed overlap between endfoot enriched transcripts, neurodevelopmental disease genes, synaptic transcripts, and PAP transcripts was significantly enriched compared to a randomized control dataset of equal size (**Fig. 3E**). Together, the shared localization of transcripts between endfeet and neural cell types, and link with neurodevelopmental disease, supports the functional relevance of endfoot enriched transcripts.

Gene ontology analysis of the 500 most enriched endfoot transcripts revealed specific functional categories (**Fig. 3F, Table S10**). The most enriched category was collagen activated tyrosine kinase receptor signaling which supports roles for endfeet in signaling and interacting with the ECM of the BM. Categories such as regulation of gene expression and mRNA splicing were also enriched. Notably, 73 of the 80 reported transcripts encoding mammalian ribosomal proteins were localized in endfeet, with 34 endfoot enriched (**Fig. 3G, Table S7**). This is consistent with the notion that endfeet are sites for local translation and consistent with observations in neurons [15, 42]. Prominent categories, including microtubule cytoskeleton organization, microtubule-based processes, and cytoskeletal organization, reinforce the importance of the local cytoskeleton in RGCs). Notably, many of these categories were also represented in the endfoot enriched proteome [18] and FMRP bound transcriptome [15]. Altogether, this rich subcellular transcriptome highlights potential new functions of transcripts for RGC endfeet.

### Specific Dynein components are subcellularly enriched in RGC basal endfeet

Microtubule cytoskeleton related transcripts were highly represented in the endfoot transcriptome. This includes components of the Dynein molecular motor (**Fig. 3H-3I**). Cytoplasmic dynein 1 (hereafter referred to as “dynein”) is involved in minus-end-directed transport of cellular components including organelles, mRNAs, proteins, and viruses [43]. Dynein is composed of a homodimer of heavy chains (HCs) and dimers of five other subunits, comprising intermediate (ICs), light intermediate (LICs), and light chains (LCs) [44]. Both *Dync1li2* and *Dynll2* were highly enriched in RGC endfeet at 730-fold and 334-fold respectively (**Fig. 3I**). In contrast, *Dync1h1* showed only 11-fold enrichment and *Dync1li1* and *Dynll1* were not enriched at all (0.26-fold and 0.46-fold; **Table S1**). Dynactin is an essential co-factor for dynein, and along with other adaptors and regulators is required for robust dynein processive motility [45]. *Dctn1,* which encodes p150(Glued), showed moderate 6-fold enrichment in endfeet. This suggests unique spatial distribution of transcripts encoding dynein components which may impact where each subunit is translated and assembled.

Given their robust enrichment in the endfoot transcriptome, we assessed the localization of *Dync1li2* and *Dynll2* in the mouse cortex. To inform potential functions of this localization, we examined *Dync1li2* localization at the pia and in the VZ/SVZ across different developmental stages (**Fig. 4A and S5A**). Using smiFISH [46], we found that *Dync1li2* and *Dynll2* strongly localized to the pial surface, with less notable localization in the proliferative zones at E14.5, E15.5, and E16.5 (**Fig. 4A and S5A**). Using RGCs labeled with EGFP by *in utero* electroporation (IUE), we determined that *Dync1li2* was especially enriched in RGC endfeet, consistent with transcriptome analysis (**Fig. 4B-4C**). In contrast to *Dync1li2* and *Dynll2* light intermediate and light chains, *Dync1h1* and *Dctn1* were uniformly distributed throughout the cortical plate and proliferative zones (**Fig. S5B-S5C**). Although *Dctn1* mRNA was not highly enriched in RGC endfeet (**Fig. 3I and S5B**), the p150 Glued protein, encoded by this transcript, localized to RGC endfeet as evidenced by immunofluorescence colocalization with EGFP-labeled endfeet (**Fig. S5D**). This validation, together with the transcriptome (**Fig. 3I**), demonstrates that transcripts encoding dynein components exhibit differential subcellular localization in RGCs. These data show that *Dync1li2* localizes strongly to RGC basal endfeet and suggest it may function across these developmental stages in endfeet.

**Fig. 4.**
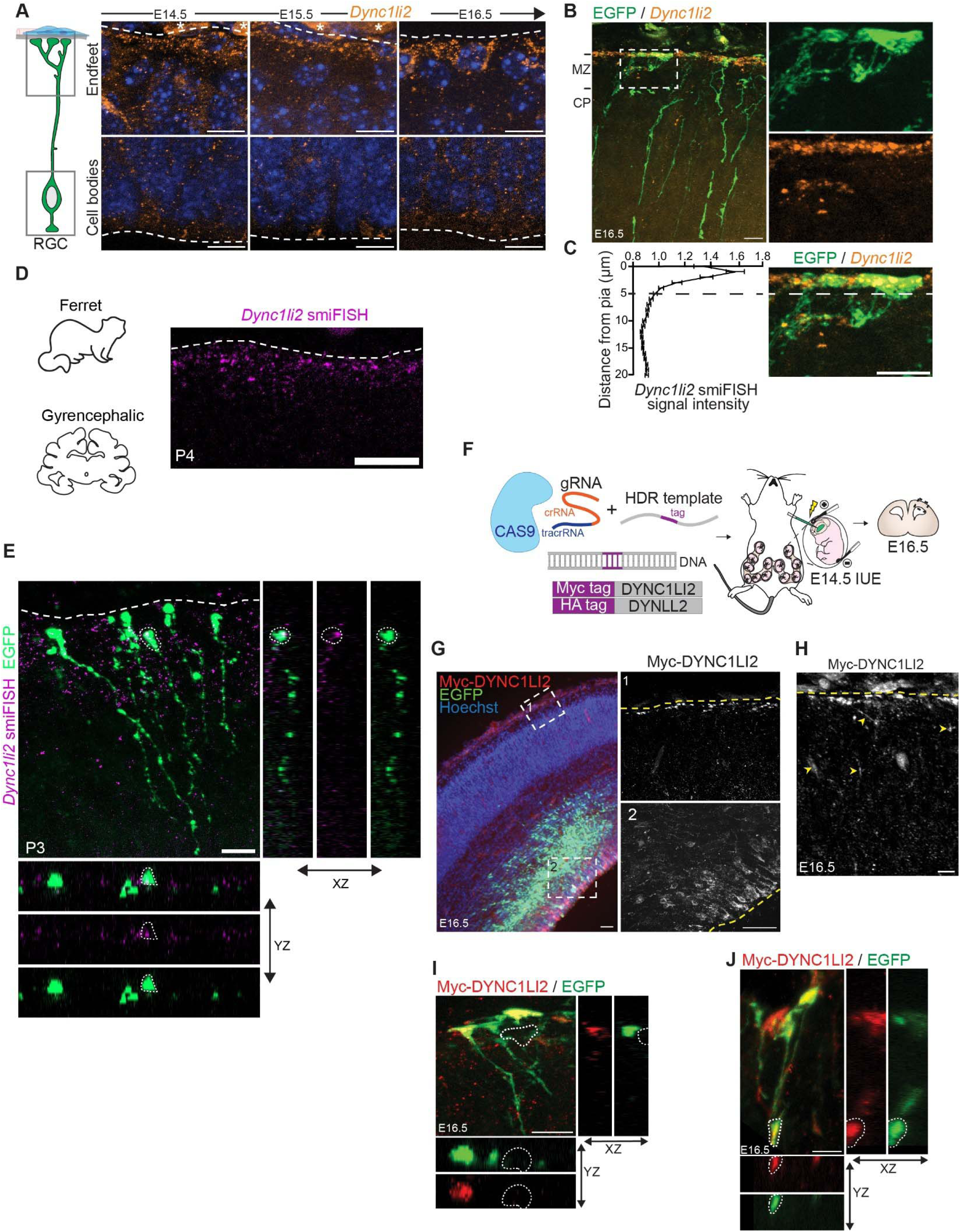
Dynein subunit *Dync1li2* mRNA and protein localize to RGC basal endfeet during cortical development. **(A)** Cartoon representation of an RGC with boxes indicating cell body and endfoot regions imaged (left). *Dync1li2* smiFISH puncta (orange) in E14.5, E15.5 and E16.5 mouse embryonic cortices with nuclei labeled with DAPI (blue) (right). White dotted lines indicate pial (top) and ventricular (bottom) borders and asterisks denote background signal from the meninges or blood vessels. **(B)** *Dync1li2* smiFISH (orange) colocalized with EGFP-electroporated RGC endfeet at the pia at E16.5 (green). White dotted line box outlines magnified area highlighted in right panels. **(C)** smiFISH signal intensity quantification from box in (B); n=12 ROIs, 4 brains from 3 litters. **(D)** *Dync1li2* smiFISH (magenta) in P4 ferret cortex imaged at the pia. White dotted line denotes pial border. **(E)** Colocalization of *Dyn1li2* smiFISH (magenta) with EGFP-electroporated RGC endfeet (green) in P3 ferret cortex. **(F)** Diagram representing the Breasi-CRISPR method used for endogenous tagging of DYNC1LI2 and DYNLL2. **(G)** Representative images showing endogenously tagged Myc-DYNC1LI2 (red) with EGFP-electroporated cells (green), and DAPI (blue) in an E16.5 cortex (left). White dotted boxes outline magnified areas highlighted in right panels. Right, Myc-DYNC1lI2 (grey) expression at the pia (top) and ventricular zone (bottom). Yellow dotted lines indicate the border between the endfeet (below) and the pia (above). **(H)** Localization of tagged Myc-DYNC1LI2 in RGC endfeet and basal processes at E16.5. Yellow dotted lines denote pial boundary. Yellow arrowheads mark tagged protein in basal processes. **(I)** Colocalization of Myc-DYNC1LI2 (red) with EGFP-electroporated RGC endfeet (green). **(J)** Localization of tagged Myc-DYNC1LI2 (red) in RGC branching points and endfeet (green). Scale bars: (A-E, G, H, I) 10µm, (J) 5 µm. Error bars: (C) SEM smiFISH, single-molecule inexpensive fluorescent in situ hybridization; RGC, radial glial cell; MZ, marginal zone; CP, cortical plate; ROIs, regions of interest; gRNA, guide RNA; crRNA, CRISPR RNA; tracrRNA, transactivating CRISPR RNA; HDR, homology-directed repair.

We next tested the conservation of *Dync1li2* localization in other species and specifically gyrencephalic mammals which have more complex basal structures [6, 47]. Towards this we performed smiFISH on ferret cortical tissue at P3 and P4, stages roughly equivalent to E15.5 in mice [48]. Ferrets have especially complex radial glial basal structures [6, 49]. *Dync1li2* signal was strong at the pial surface and co-localized with basal endfeet labeled by electroporated EGFP (**Fig. 4D-4E**). These data demonstrate that *Dync1li2* subcellular distribution is conserved in ferret RGCs, reinforcing its potential functional importance in endfeet.

We next examined localization of DYNC1LI2 and DYNLL2 proteins in RGCs. For this, we used Breasi-CRISPR, a genome editing IUE method to tag endogenous DYNC1LI2 and DYNLL2 proteins with Myc and HA, respectively [50, 51] (**Fig. 4F**). Both tagged proteins localized to the cell soma and along presumptive basal processes and branches (**Fig. 4G, 4H and S5E**). Notably, Myc-DYNC1LI2 overlapped with EGFP-labeled endfeet and localized to RGC branch points (**Figs. 4I-4J**). This localization pattern further suggests DYNC1LI2 may function in RGC basal structures. Thus, both RNA and protein for DYNC1LI2 and DYNLL2 strongly localize to RGC basal endfeet suggesting local translation and function of subcellular *Dync1li2* and *Dynll2*.

### *Dync1li2* is critical for RGC basal branching and non-cell autonomous regulation of interneuron number in the MZ

Dynein components are essential for formation of branches in neurons [52, 53]. Given this known function and the significant enrichment and localization of DYNC1LI2 in RGC basal structures, we postulated this dynein subunit may also influence morphology of RGCs during cortical development. To test whether *Dync1li2* influences cell morphology, we introduced either scrambled (control) or *Dync1li2* siRNAs into E15.5 brains by IUE, along with a membrane-bound EGFP driven by an RGC-specific promoter (pGLAST-EGFP-CAAX) for sparse labelling of RGCs (**Fig. 5A**). Following 24h knockdown (KD), *Dync1li2* levels were significantly reduced in the *Dync1li2* siRNA condition compared to control (**Fig. 5B-5C**). *Dync1li2* depletion was especially evident close to the pia where endfeet reside (**Fig. 5B-5C and S5F-S5H**). We also observed a discrete reduction in RGC cell bodies, consistent with low levels of expression in this compartment (**Fig. S5F-S5H**). Sorting of RGC endfeet further supported that siRNA led to significant reduction of *Dync1li2* (**Fig. S6A**).

**Fig. 5.**
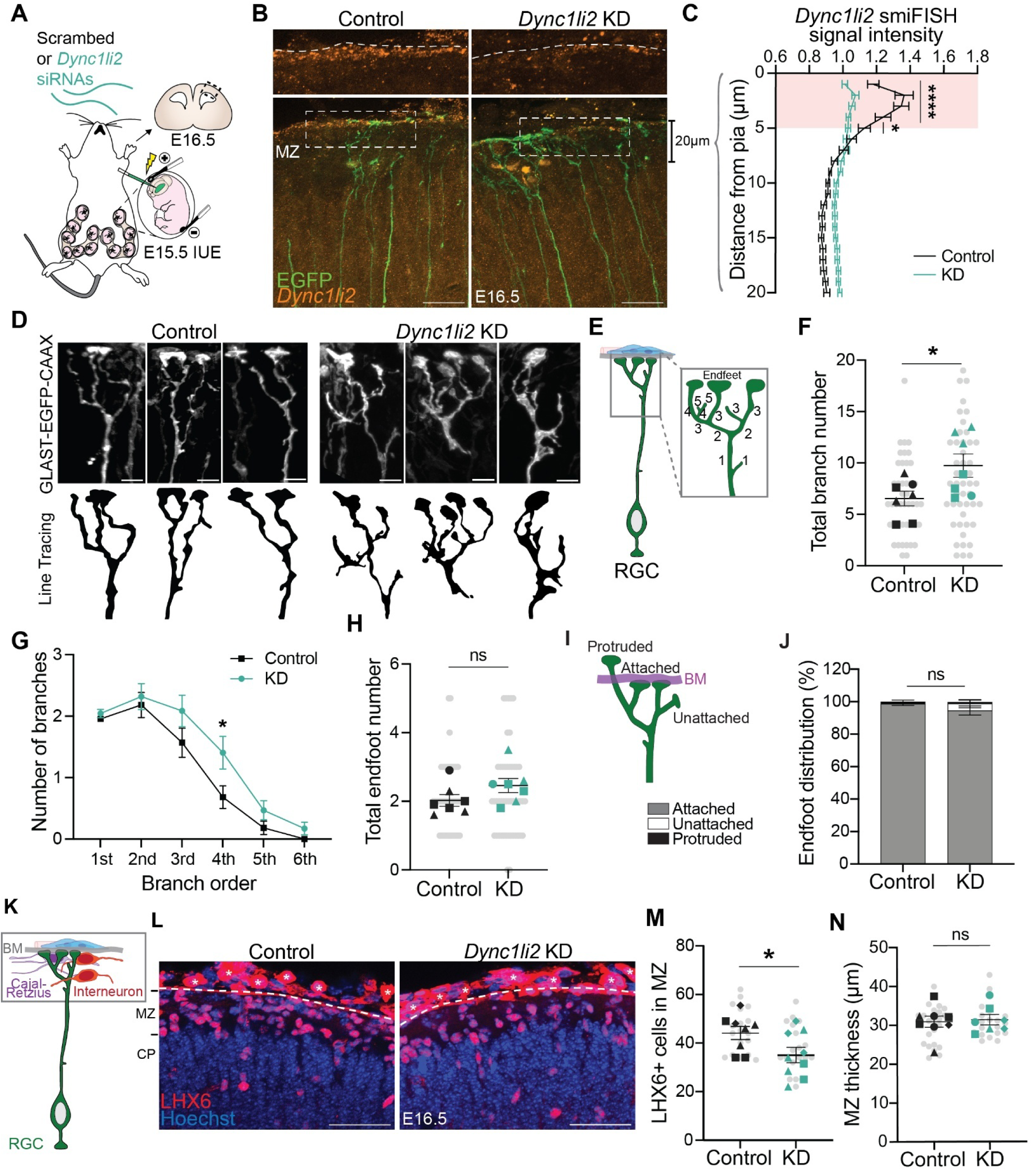
*Dync1li2* siRNA knockdown increases RGC basal complexity and non-cell autonomously impacts interneuron number in the MZ. **(A)** Cartoon outlining the approach used for siRNA knockdown *Dync1li2* transcripts in mouse embryonic brains. **(B)** Representative images depicting *Dync1li2* smiFISH (orange) in EGFP-electroporated regions (RGCs in green) at E16.5. White dotted line boxes outline magnified area in the top panels, highlighting *Dync1li2* mRNA expression at the pia in control (left) and KD (right) cortices. **(C)** smiFISH signal intensity quantification comparison between control (black) and *Dync1li2* KD (teal) samples in the first 20µm of the MZ. Pink box denotes first 5μm of the MZ where endfeet reside. n=14 ROIs from 7 control brains and n=19 ROIs from 10 KD brains, from 3 litters **(D)** Representative images of 3D reconstructed control and *Dync1li2* KD RGCs labeled with GLAST-EGFP-CAAX. Fluorescent images (top), and line tracing (bottom) of individual RGC basal processes and endfeet. **(E)** Schematic detailing branch number and order quantification criteria for panels E-G. **(F)** Quantification of total branch number per RGC. **(G)** Quantification of number of branches per branch order. **(H)** Quantification of total endfoot number per RGC. **(I)** Schematic detailing quantification criteria for endfoot position relative to the BM. **(J)** Quantification of endfoot position represented as % of total endfeet analyzed. (F-H, J) n=44 cells from 7 control brains and n=46 cells from 7 KD brains, from 3 litters. **(K)** Cartoon representation of an RGC (green) and the local niche of basal endfeet, which include the basement membrane (BM, grey), fibroblasts (blue), vasculature (pink), Cajal-Retzius (CR) cells (purple), and interneurons (red). **(L)** Representative images of LHX6 staining (red) of interneurons and Hoechst (blue) at E16.5. White dotted lines mark boundary between pia and MZ. Black asterisks denote background signal from the meninges or blood vessels. **(M)** Quantification of LHX6+ cells in the MZ. n=18 sections, 8 control brains, and n=19 sections 9 KD brains, from 3 litters. **(N)** Quantification of MZ thickness. n=24 measurements from 8 control brains, and n=20 measurements from 7 KD brains, from 4 litters. Scale bars: (B) 20µm, (D) 5µm, (L) 50µm; Stats: (C, G) Two-way ANOVA with Bonferroni’s multiple comparisons test; (F, H, M, N) Student unpaired, two-tailed t test; (J) Two-way ANOVA with Sidak’s multiple comparisons test. Error bars: SEM (C, F, G, H, J), SD (M, N). Super-plots depict ROIs or cells in gray dots and individual data representing different brains in black (control) or teal (KD) dots; litters are coded by shape. ns p-value > 0.05, *p-value < 0.05, ****p-value < 0.0001. siRNAs, small interfering RNAs; IUE, in utero electroporation; MZ, marginal zone; CP, cortical plate; KD, knockdown; RGCs, radial glial cells; ROIs, regions of interest; BM, basement membrane.

Having established a paradigm to deplete *Dync1li2* in RGCs, we next assessed the functional impact on RGC morphology. Towards this we generated 3D reconstructions of individual RGCs and measured morphological features including branch number, branching location and endfoot number (**Fig. 5D**). First, we quantified the total branch number density according to their order assignments, as previously described (**Fig. 5E**) [18, 19]. We observed a significant increase in total branch number in *Dync1li2*-depleted RGCs compared with controls (**Fig. 5F**). This was driven by a significant increase in the number of higher order branches in *Dync1li2* KD cells (**Fig. 5G**). However, this increased RGC complexity had no impact on the location of RGC branching relative to the pia (**Fig. S6B-S6C**). Both control and *Dync1li2* KD showed similar number of endfeet per cell (**Fig. 5H**). There were no changes in the position of endfeet relative to the BM, as assessed by Laminin staining, with the vast majority properly attached to the BM (98% and 94%; **Fig. 5I, J, S6D**). Thus, loss of *Dync1li2* induces aberrant formation of RGC branches, however these branches do not result in new endfeet. Altogether, these results indicate *Dync1li2* is critical for morphological complexity of RGC basal structures.

To examine the consequences of increased branching complexity on cortical architecture, we assessed cells populating the MZ (**Fig. 5K**). We previously determined that alterations in RGC integrity and branching complexity impact interneuron number and position in the MZ [18, 19]. In particular, absence of RGC endfeet, in a *Myh10* mutant, led to more interneurons in the MZ [18]. Thus, we hypothesized that in *Dync1li2* mutants, which have more complex RGC basal structures, interneurons may be depleted. Indeed, quantification of LHX6+ cells showed a significant decrease in the number of interneurons in the MZ of *Dync1li2* knockdown cortices compared with controls (**Fig. 5L-5M**). Importantly, there was no notable impact on MZ thickness indicating this is not simply due to loss of this zone (**Fig. 5N**). Likewise, there was no change in the number of P73+ Cajal-Retzius cells (CR cells) in the MZ (**Fig. S6E-S6F**). This aligns with previous findings showing that at these stages, RGC endfeet do not grossly impact CR number [18, 19]. It is also consistent with the earlier timing of CR neuron migration, which may be completed before *Dync1li2* knockdown [54]. In sum, these data indicate that more complex RGC basal structures impair interneuron number in the MZ.

### LOCAL-KD: an *in vivo* tool for subcellular gene knockdown

The *Dync1li2* knockdown experiments demonstrate a role for DYNC1LI2 in controlling RGC morphology, however they raise the important question as to whether this is due to a local requirement in endfeet. Because transcripts enriched in endfeet are also expressed in the cell body, depletion by siRNA removes the RNA of interest from both cellular compartments. In previous studies of endfoot localized *Arhgap11a*, our group determined local functions using rescue experiments with specific versions of the transcript that localize to endfeet [19]. However, this approach relies on overexpression and requires knowing the localization element for each gene of interest. Thus, rescue approaches are technically challenging, labor intensive and not easily scaled.

Hence, we developed a new method, <u>Lo</u>cal <u>Ca</u>s13d mRNA <u>L</u>evel <u>K</u>nock<u>d</u>own (LOCAL-KD), to specifically deplete mRNAs of interest, including *Dync1li2*, in subcellular compartments of RGCs. Towards this, we used the CRISPR-Cas13 method [55, 56]. Cas13 is an RNA nuclease which targets specific mRNAs using gene specific guide RNAs (gRNAs). We hypothesized that by controlling the localization of the Cas13 protein we could achieve spatially targeted gene knockdown. Amongst the many versions of Cas13, we selected Cas13d as it is functional in cell culture and *in vivo*, including zebrafish and mice, with limited off target effects [57–63]. Additionally, it is significantly smaller than other Cas13 variants allowing for easier delivery by IUE [58]. However, its functionality in the developing mouse cortex has yet to be reported and it has not been used previously for subcellular manipulation.

To produce compartmentalized knockdown in either the basal endfeet or cell body we needed to differentially localize the Cas13d machinery to each subcellular compartment (**Fig. 6A**). We first optimized promoters for this approach. We introduced a previously published nuclear localizing Cas13d (NLS-Cas13d-NLS-HA-T2A-EGFP) under control of either a U6 or pCAG promoter into embryonic cortices by IUE [59]. After 24 hours, the U6 promoter drove only low levels of EGFP expression, while the CAG promoter led to higher expression relative to mCherry control (**Fig. S7A**). Next, we sought to direct Cas13d to distinct sub-compartments of RGCs. Overexpression can cause proteins that do not normally localize to endfeet such as EGFP or mCherry to diffuse throughout the entire cell including the endfeet. To retain Cas13d in the cell body, we used two nuclear localization sequences (NLS) on Cas13d as previously [59] (**Fig. 6A**). We IUE’d the pCAG-NLS-Cas13d-NLS-HA-T2A-EGFP plasmid into embryonic cortices at E15.5 along with pCAG-EGFP to label RGCs including endfeet. After one day, we stained for HA and performed smiFISH against Cas13 to measure both NLS-Cas13d protein and mRNA localization, respectively, within the RGCs. Neither NLS-Cas13d protein nor *Cas13d* RNA localized in EGFP labeled endfeet (**Fig. 6B-6C**). However, as expected, we observed *Cas13d* mRNA and protein in the cell bodies where it is transcribed (**Fig. 6B**). Therefore, the NLS was sufficient to prevent passive localization of the Cas13d to endfeet, after 24 hours.

**Fig. 6.**
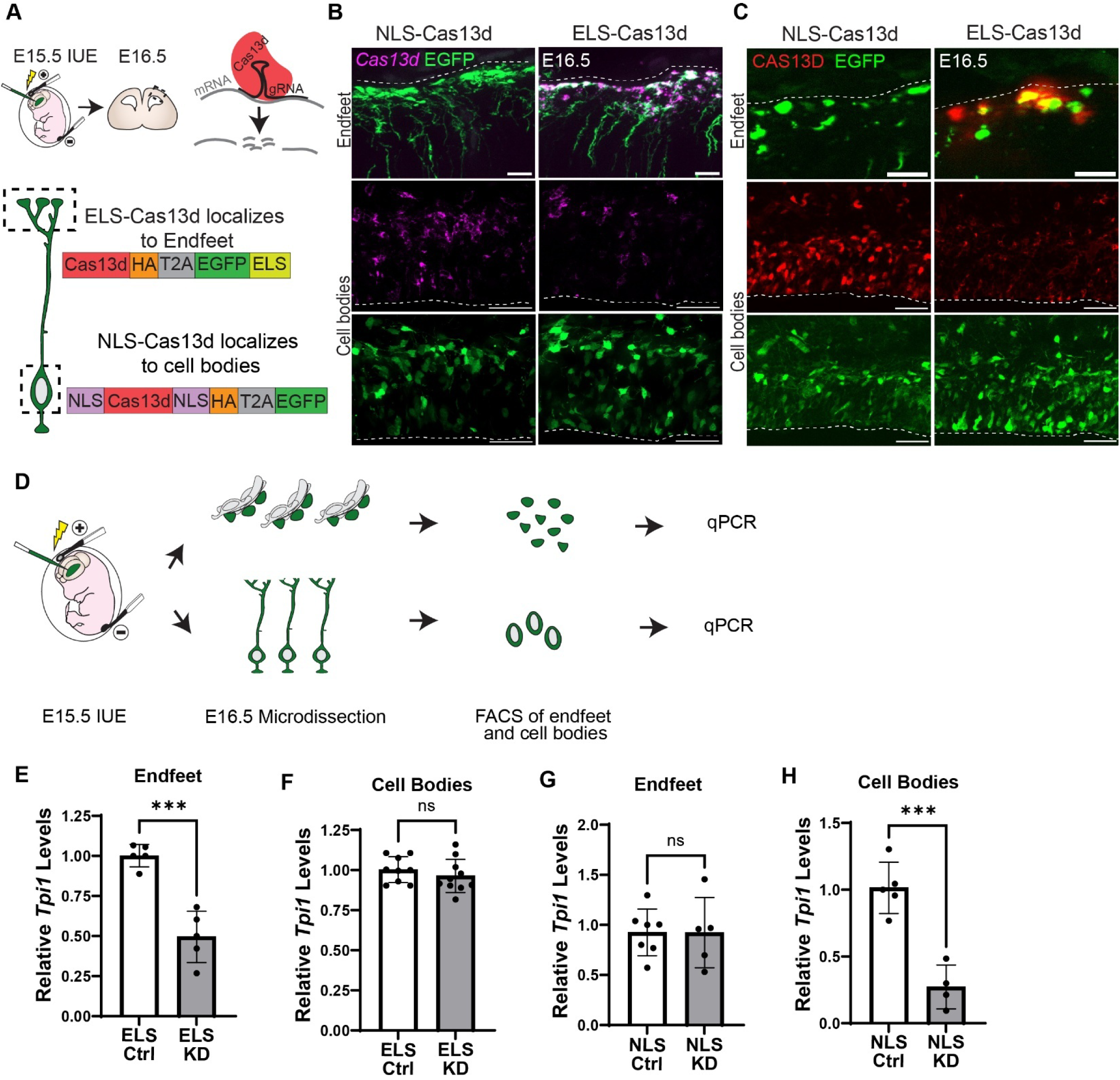
Development of the LOCAL-KD system for degrading transcripts in subcellular compartments *in vivo*. **(A)** Cartoon representation of experimental paradigm and Cas13d system. Expression by IUE at E15.5 with analysis at E16.5. **(B, C)** Expression of NLS-Cas13d (left) and ELS-Cas13d (right) in E16.5 cortices with smiFISH against *Cas13d* mRNA (magenta) (B) and immunofluorescence against HA labeling Cas13d-HA protein (red) (C). EGFP denotes electroporated endfeet (top) and cell bodies (bottom). White dotted lines mark the ventricular and pial boundaries. **(D)** Cartoon representation of experimental design used to test subcellular knockdown of *Tpi1 in vivo* (quantified E-H) **(E)** Relative levels of *Tpi1* in endfeet following electroporation with ELS-Cas13d. n=5, 4-7 brains pooled per replicate. **(F)** Relative levels of *Tpi1* in cell bodies following electroporation with ELS-Cas13d. n=9 Ctrl, 10 KD, 3 brains pooled per replicate. **(G)** Relative levels of *Tpi1* in endfeet following electroporation with NLS-Cas13d. n=7 ctrl, 5 KD, 8-12 brains pooled per replicate. **(H)** Relative levels of *Tpi1* in cell bodies following electroporation with NLS-Cas13d. n=5 ctrl, 4 KD, 3 brains pooled per replicate. Scale bars: (B-C) 10μm endfeet (top), 50μm cell bodies (bottom). (E-H) points represent biological replicates. Stats: (E-J) Student’s unpaired, two-tailed t test. Error bars: (E-J) SD. ns p-value > 0.05, *p-value < 0.05, ***p-value < 0.001.

To drive the localization of Cas13d to endfeet we replaced the NLS motifs with the 5′ UTR of *Arhgap11a*, which has been previously shown to localize mRNA to endfeet [19], referred to as the endfoot localization sequence (ELS). We did not add localization elements to the gRNAs as they are small (∼60nt) and their function may be impaired by this addition (∼500nt). One day following electroporation of the pCAG-Cas13d-HA-T2A-EGFP-ELS plasmid with pCAG-EGFP, we observed strong localization of both *Cas13d* mRNA and protein to endfeet (**Fig. 6B-6C**). While ELS-Cas13d mRNA was present in the cell body where it is transcribed, the protein was barely detectable in the cell body. This demonstrates that the ELS drives specific localization of *Cas13d* mRNA to endfeet, where it is presumably locally translated. In sum, this approach enables us to differentially localize Cas13d to cell bodies or endfeet.

We next tested the ability of these localized forms of Cas13d to target RNAs in different compartments. From these localization data we predicted the NLS-Cas13d would deplete target genes in the cell body with little knockdown in the endfeet and the ELS-Cas13d would specifically deplete targets in the endfeet with little knockdown in the cell body. To test this, we identified a proof-of-concept transcript to target with ELS-Cas13d and NLS-Cas13d. We leveraged our subcellular transcriptome to identify mRNAs that were moderately expressed in both cell bodies and endfeet, and that showed specific expression in RGCs and negligible expression in surrounding cells. Genes expressed in Neuro2A cells were prioritized to allow for an *in vitro* model for screening gRNAs. *Tpi1* met all selection criteria with FPKMs of 66.4 and 204.8 in the cortex and endfeet respectively, undetectable expression in E14.5 meninges, and expression in N2A cells (**Fig. S7C)** [23, 64]. Single cell RNA sequencing data also showed enriched expression in RGCs compared to neurons across development [65] **(Fig. S7B)**. *Tpi1* expression in RGC cell bodies and endfeet was validated by smiFISH (**Fig. S7D, E**). gRNAs for *Tpi1* were designed and validated to KD *Tpi1* using NLS-Cas13d in N2A cells 24 hours post transfection (**Fig. S7F**). This provided a proof-of-principle transcript for testing localized knockdown.

We next tested the specificity of spatial targeting with ELS-Cas13 and NLS-Cas13 *in vivo* by introducing the Cas13d and gRNA by IUE, collecting endfeet and cell bodies by FACS after 24 hours, followed by qPCR to measure mRNA levels (**Fig. 6D**). Using ELS-Cas13 and *Tpi1* gRNA, *Tpi1* levels were reduced 51% in basal endfeet compared to ELS-Cas13 control (**Fig. 6E**). Cell bodies collected from the same brains showed no difference in *Tpi1* levels between conditions (**Fig. 6F**). This revealed that ELS-Cas13 causes mRNA knockdown specifically in the endfeet without impacting expression in cell bodies. In contrast, NLS-Cas13 showed a 73% reduction in cell bodies without knockdown in endfeet (**Fig. 6G-6H**). Therefore, NLS-Cas13 can produce a cell body specific depletion of *Tpi1* without changing *Tpi1* levels in endfeet. These data validate LOCAL-KD for targeting mRNAs in subcellular compartments of RGCs.

### Subcellular *Dync1li2* is required for RGC basal morphology

Leveraging our new LOCAL-KD method, we next established tools to assess the subcellular requirements of *Dync1li2* in RGCs. gRNAs specific for *Dync1li2* were designed and validated in N2A cells (**Fig. S7G**). The gRNA with the most efficient knockdown was also validated *in vivo* by qPCR of purified endfeet (**Fig. 7A**). 24 hours post introduction of ELS-Cas13d and *Dync1li2* gRNA resulted in a 62% reduction in endfoot localized *Dync1li2* compared to control (**Fig. 7B**). In contrast, from these same brains, these conditions did not produce any significant knockdown in the cell bodies (**Fig. 7C**). In both control and KD cell bodies, *Dync1li2* levels were variable and low as evidenced by RNA sequencing, smiFISH, and qPCR (**Supplemental Table 1, Fig. S5F-S5H, 7C**). Therefore, with this method we successfully produced endfoot targeted knockdown of *Dync1li2 in vivo*.

**Fig. 7.**
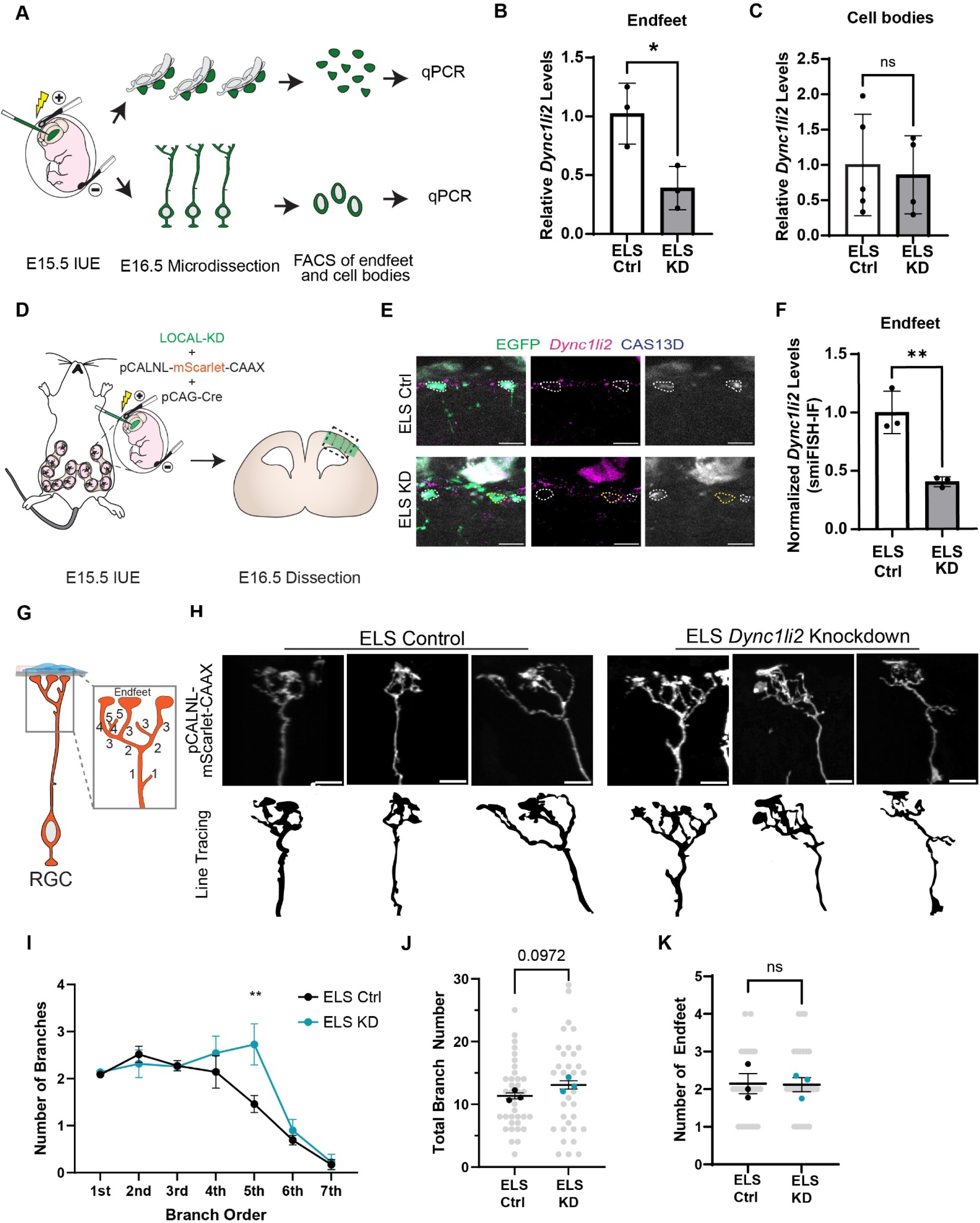
LOCAL-KD shows subcellular expression of *Dync1li2* is required for RGC basal morphology. **(A)** Cartoon representation of experimental design used to test subcellular knockdown of *Dync1li2 in vivo* **(B)** qPCR of *Dync1li2* levels in purified endfeet after IUE with ELS-Cas13d. n=3, 3-4 brains pooled per replicate. **(C)** qPCR of *Dync1li2* levels in sorted cell bodies after IUE with ELS-Cas13d. n= 4-5, 1-2 brains pooled per replicate **(D)** Cartoon summary of experimental paradigm. **(E)** smiFISH-IF for *Dync1li2* mRNA (magenta) and ELS-CAS13 protein (blue) in EGFP+ endfeet (green). White dotted circles denote CAS13 positive endfeet. Yellow dotted circle marks an example of an EGFP+, CAS13-endfoot. **(F)** Quantification of *Dync1li2* mRNA in CAS13D positive endfeet by smiFISH-IF. Points represent brains. Sections normalized to *Dync1li2* mRNA levels in CAS13D negative endfeet from the same brain. Brains KD normalized to Ctrl condition. n=3 per condition 2-4 sections quantified per brain from 1 litter. **(G)** Schematic detailing branch number and order quantification criteria **(H)** Representative images of 3D reconstructed control and *Dync1li2* KD RGCs labeled with pCALNL-mScarlet-CAAX. Fluorescent images (top), and line tracing (bottom) of individual RGC basal processes and endfeet. **(I)** Quantification of branch complexity. **(J)** Quantification of branch number. **(K)** Quantification of endfoot number. (I-K) n= 36 Ctrl cells and 35 ELS-KD cells from 3 brains per condition Scale Bars: (E, H) 10μm. Stats: (B, C, J, K) Student’s unpaired, two-tailed t test. (I) Two-way ANOVA with Bonferroni’s multiple comparisons test. Error bars: (B, C, F) SD (I-K) SEM. Super-plots depict cells in gray dots and individual data representing different brains in black (control) or teal (KD) dots. ns p-value > 0.05, **p-value < 0.01.

We next used this paradigm to assess the impact of localized depletion of *Dync1li2* upon the morphology of RGC basal structures. Towards this we IUE’d LOCAL-KD tools along with sparse Cre and a Cre-inducible membrane-bound mScarlet (**Fig. 7D**). This approach enabled us to sparsely label control and *Dync1li2* endfoot knockdown RGCs for detailed morphology analysis. Successful knockdown of *Dync1li2,* in sparsely labeled endfeet, was validated by smiFISH-IF. A significant 60% decrease in endfoot localized *Dync1li2* was observed in CAS13D positive ELS KD endfeet compared to ELS control (**Fig. 7E-7F**). Using the same samples in which *Dync1li2* depletion was validated, 3D reconstructions were rendered and morphology analysis performed as previously with the *Dync1li2* siRNA samples (**Fig. 7G-7H**). Knockdown of *Dync1li2* in RGC endfeet led to significantly increased complexity, as reflected by branch order (5^th^ order) (**Fig. 7I**). ELS-KD RGCs also trended towards (p=0.097) increased total branches per cell (**Fig. 7J**). ELS-Ctrl and ELS-KD RGCs had equivalent numbers of endfeet per cell (**Fig. 7K**). Therefore, these results largely phenocopy our findings with siRNA knockdown, which robustly depleted *Dync1li2* from the entire RGC. Altogether, these data using LOCAL-KD demonstrate that endfoot *Dync1li2* is locally required for RGC morphology.

## Discussion

RNA localization is a fundamental feature across organisms and cells, including polarized radial glial progenitors of the developing brain. Two key questions are: what is the breadth of mRNA localization in RGCs and how does it influence cortical development? Tackling both questions requires a comprehensive understanding of subcellular transcriptomes, as well as experimental paradigms to study RNAs in a subcellular fashion. In this work we developed new methods to purify RGC subcellular compartments, which allowed us to discover a rich, subcellular transcriptome of RGC, far from the cell soma. Our discoveries generate new hypotheses about endfoot function in the developing cortex beyond their physical roles in controlling neuronal migration. Functional interrogation of one highly enriched transcript revealed essential requirements of the Dynein complex component *Dync1li2* in restricting basal branching complexity. We further develop LOCAL-KD for spatially targeted mRNA knockdown using Cas13d, which we use to show that *Dync1li2* is required sub-cellularly in endfeet. Altogether, our study opens up a new understanding of RGC subcellular structures and new layers of gene regulation at play in the developing brain (**Fig. 8**). More broadly, we discover foundational insights and establish new technical innovations for understanding subcellular RNA localization in the nervous system.

**Fig. 8.**
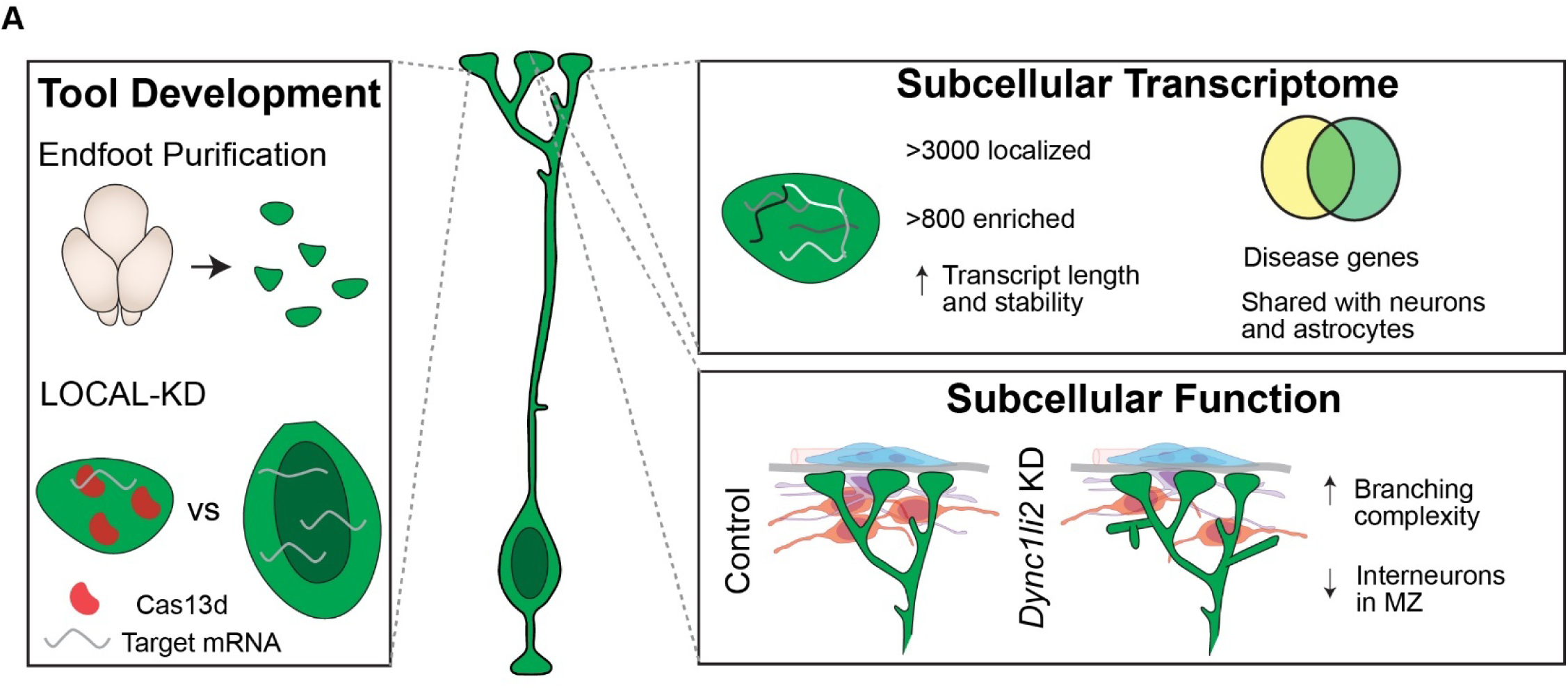
(A) Cartoon model of the main findings of the study. This study generates new experimental paradigms to purify RGC endfeet and manipulate subcellular transcriptomes. We use these new tools to define the subcellular transcriptome of RGCs and to demonstrate that Dynein components are critical for morphology of RGC basal structures and cortical development.

### Transcriptomes of purified endfeet reveal a new layer of subcellular gene expression in radial glia

We discovered over 3,000 unique transcripts in RGC basal endfeet, 815 of which are highly enriched and abundant compared to cell bodies. This dramatically expands the list of localized RNAs from several hundred to several thousand [15–17, 19, 21]. Our transcriptome suggests that RGC basal endfeet may be more multi-functional than previously appreciated. Endfoot enriched transcripts include those with known functions in signal transduction and endocytosis, supporting putative roles for endfeet in signaling. It also includes collagens, laminins, and fibrillins suggesting that endfeet could be sites of subcellular ECM production. Consistent with this, many of these ECM associated RNAs were also represented in the endfoot proteome [18]. Surprisingly, far from the nucleus, transcripts encoding DNA binding regulators were also enriched. Some of these may function non-canonically, as similar categories of transcripts are found in axons [66, 67]. Alternatively, these transcripts could be sequestered in the endfeet and trafficked back to the cell body as needed, as has been proposed for cell cycle regulators [16, 68]. Together this new transcriptome underlies potential new functions for endfeet in intracellular and extracellular signaling, generating new testable hypotheses for future investigation.

How are these mRNAs trafficked to, localized, and controlled within endfeet? FMRP was previously shown to be critical for mRNA trafficking and localization in endfeet. Indeed, FMRP-bound transcripts [15] were significantly enriched in our transcriptome. However, these represent only a fraction of the transcriptome, suggesting that FMRP is just one of many regulatory RNA binding proteins. In future experiments, it will be valuable to uncover the mRNA targets of additional RNA binding proteins in endfeet and their role in different aspects of mRNA metabolism in endfeet. This may reveal the extent to which RNA binding proteins control functionally discrete groups of transcripts as well as how they spatially and temporally control RNAs in RGCs.

In many systems, including RGC endfeet, localized RNAs are competent to undergo translation [15]. Compared to the endfoot proteome, about half of the proteins were present at the mRNA level [18]. Notably this proteome is unlikely comprehensive as it relied on overexpression of a cytoplasmic BioID. Nevertheless, we predict this subset of overlapping transcripts may be locally translated in RGC endfeet, while the remainder may be produced in the cell body and trafficked to endfeet. These distinct mechanisms could be related to transcript size as endfoot localized transcripts are significantly longer than those in the cell body. Indeed, it may be more energetically favorable to transport mRNA and locally translate larger proteins [69]. Interestingly, components of the ubiquitin-proteasome pathway are also enriched in endfeet at both the RNA and protein level. This suggests that local protein degradation may reciprocally be important for gene regulation in endfeet.

Our study raises additional fascinating questions about RGC subcellular composition and function. In this study, we measured the subcellular transcriptome at a single stage (E15.5), yet tight spatiotemporal control is likely important and even distinct over the course of cortical development. Indeed, we previously observed that localization of some mRNAs, such as *Myh9* and *Myh10,* can dynamically change over the course of development [18]. Throughout development, RGC morphology becomes more elaborate and RGC function evolves as they make distinct excitatory neurons and then glia. Our experimental paradigm can thus be used to measure the subcellular RGC transcriptome composition at diverse stages to further understand RGC biology and cortical development. Additionally, the ability to isolate pure populations of endfeet makes it possible to carry out multi-omics analyses to profile proteome, RNA modifications, and diverse RNA types (cirRNA, tRNA etc). Likewise, our study and previous studies [19] reveal some transcripts have evolutionarily conserved localization to endfeet, as we have shown for *Dync1li2* localization to ferrets. However, the extent to which RGC endfoot localization is widespread across species remains unknown. Further, primates and gyrencephalic species have abundant oRGs with basal endfeet, making it important to understand the composition of cell-specific endfeet. Spatial approaches may also be valuable to define the composition of individual endfeet and endfoot heterogeneity. Altogether, this new endfoot transcriptome reinforces previous findings while also suggesting exciting new roles for local gene regulation in endfeet.

### Endfoot localized *Dync1li2* controls complexity of RGC basal structures and cortical organization

Strikingly, we discovered that many endfoot-enriched transcripts encode motor proteins, including both dyneins and kinesins. Our study highlights new functions for *Dync1li2* in cortical development. Dynein components have been implicated in cortical development including interkinetic nuclear migration and mitotic entry during RGC proliferation [70–75], as well as neurodevelopmental disease [76–79]. We discover a new function for *Dync1li2* in RGC branching complexity and interneuron organization in the MZ. This complements observations with other localized endfoot transcripts, showing that absence of RGC endfeet and branches in the MZ increases interneurons [18] and that reduced basal branching complexity affects interneuron organization [18, 19]. Importantly, our data suggest that *Dync1li2* controls interneuron density by acting non-cell autonomously in RGCs, since the IUE delivered siRNAs to RGCs in the VZ, but not interneurons. Together these data support a model in which RGC basal structures influence interneuron position, which may be mediated by physical and/or chemical interactions.

Endfeet have a unique composition of dynein components compared to the cell body, raising the interesting notion that compartment specific complexes may exist. While *Dync1li2* and *Dynll2* are highly enriched in endfeet at the RNA level, the heavy chain is not. Instead, DHC1 protein may be transported from the cell body to endfeet where the dynein complex forms together with locally translated components. It is also possible that low levels of *Dhc1* transcript may be locally translated in endfeet. Moreover, in contrast to *Dync1li2* localization to endfeet, the paralog *Dync1li1* showed biased localization to the cell body (Table S1). Differential composition of dynein complexes has been shown to enable functional versatility, regulating cargo selectivity and motor properties [43, 80]. Thus, DYNC1LI1 and DYNC1LI2 may define distinct cytoplasmic dynein populations with unique cargoes in each compartment [81, 82]. Of note, DYNC1LI1 and DYNC1LI2 have unique roles during cortical development, with only the former required for RGC INM and multipolar-to-bipolar transition of migrating neurons [70–74]. Future studies may address the fascinating question as to how Dynein controls RGC morphology including the identities of its local cargoes and its subcellular activity. More broadly, Dynein models a paradigm for other multimeric protein complexes where subcellular functions are dictated by compartment specific expression of components.

### LOCAL-KD is a new tool for further exploration of local transcript function

Technical innovation has led to increased subcellular profiling of a wide variety of cell types. However, tools to interrogate the local functions of these genes are lacking. Our study addresses this gap by developing a new tool, LOCAL-KD, for spatially regulated targeting of mRNAs *in vivo*. We leverage this tool to demonstrate that the endfoot localized *Dync1li2* is required to curb formation of high order, complex branches. While local *Dync1li2* depletion mirrors the complexity and endfoot number seen with whole cell siRNA mediated knockdown, the observed increase in overall branch number is less robust. siRNA led to a more pronounced reduction of *Dync1li2* compared to LOCAL-KD which may explain this phenotypic difference. This may be due to technical differences. LOCAL-KD requires the *Cas13d* mRNA to be transported to endfeet and locally translated before it is active. Therefore, even though we assayed samples at an endpoint of 24 hours post-IUE with both paradigms, the actual duration over which *Dync1li2* is depleted may be less in the LOCAL-KD condition, causing a milder phenotype. Further, although we focused our analysis on only cells with Cas13d positive endfeet, some quantified cells may have weak knockdown of *Dync1li2*. For transcripts like *Dync1li2* which have especially high levels in endfeet, robust Cas13d expression may be necessary.

These differences in siRNA and LOCAL-KD phenotypes may inform biological mechanisms by which Dynein controls branching. For example, *Dync1li2* transcripts are localized to RGC branch points, which may contribute to branching regulation, as in neurons [83]. Although we expect siRNAs would target mRNAs localized to branch points, LOCAL-KD may be more restricted to endfeet. Moreover, stronger branching phenotypes may result from collective contributions of many adjacent mutant RGCs. Such a result could occur with siRNA electroporation which causes more widespread knockdown of many RGCs [19], compared to LOCAL-KD targets.

LOCAL-KD may be employed with different variations in the cortex and beyond. While we demonstrate its use for targeting individual transcripts, multiplexed gRNAs or gRNA libraries could be used to simultaneously screen multiple localized transcripts. Additionally, the Cas13d component could be replaced with other CRISPR enzymes, such as mini, high fidelity, and photoconvertible versions of Cas13 [84–86]. The ability to edit, excise, endogenously tag, or alternatively splice localized mRNAs will greatly expand the toolkit for understanding RNA localization and subcellular function in RGCs and other cell types [59, 87–91]. We employed a localization element from the 5′ UTR of *Arhgap11a* which is also subcellularly localized in neurons and astrocytes [30, 32]. Therefore, these cell types may require minimal modifications to the method if the *Arhgap11a* localization element is conserved. In other models, customization of the localization element in the Cas13d vector should allow for local knockdown in the sub-compartment and cell type of choice.

## Summary

This study provides foundational data and methods for understanding local gene regulation in RGCs and throughout the nervous system. Our endfoot purification method will allow for future studies expanding our understanding of endfoot composition and function with diverse omics approaches, developmental stages, and species. The E15.5 subcellular transcriptome detailed in this work reveals a wealth of targets for further exploration including disease associated genes and those with shared subcellular localization in neurons and glia. Future functional studies will be facilitated by our newly developed method for subcellular knockdown *in vivo*. In sum, our study reveals important new principles about gene regulation in RGCs, and mRNA localization and translation in an *in vivo* developing nervous system.

## Materials and methods

### Mouse husbandry

All animal use was approved by the Duke Institutional Animal Care and Use Committee on 4/1/22 under approval number A060-22-03. The following mouse lines were used: *Nestin*-EGFP (Gift, Qiang Lu) [22] and wild type C57BL/6J mice (Jackson Laboratory) for transcriptomics and validations. Additional experiments were performed on CD1 mice (Charles River Laboratories). Background: *Nestin-*EGFP C57BL/6J. Embryonic stage E0.5 defined as the morning the plug was identified.

### Statistical methods and rigor

Description of specific n numbers, statistical tests, and p-values for each experiment are reported in the Fig. legends. For all experiments, male and female mice were used and littermates were compared when possible. All analyses were performed by one or more investigators blinded to condition.

### Fluorescence-activated cell sorting (FACS)

For RNA sequencing, *Nestin*-EGFP positive embryos were identified by EGFP fluorescence at the time of dissection. Samples were collected at E15.5. Endfeet were microdissected from the cortex by peeling the meninges and BM. Endfoot peels and the remaining cortices were separately pooled in cold Diethyl pyrocarbonate (DEPC) PBS and kept on ice until all samples were collected. Samples were centrifuged at 300g for 5min to pellet. Supernatant was discarded. 250µl prewarmed 0.25% trypsin with EDTA was added to each sample. Samples were incubated in a 37°C water bath for 10min agitating once after 5min. 750µl of 1x trypsin inhibitor in cold HBSS was added to each sample. Samples were dissociated by gently pipetting, centrifuged at 300g for 5min, and supernatant was removed. Samples were resuspended in cold HBSS and passed through a 30µm cell-strainer. Propidium iodide was added 1:1,000 to the endfoot samples. Endfeet and cell bodies were sorted at 6°C using a B-C Astrios cell sorter and 70µm nozzle with gating for forward scatter, side scatter, propidium iodide, and EGFP. Samples were sorted directly into Trizol LS (Thermo Fisher, 10296010) and stored at -80°C until RNA extraction.

### RNA extraction, cDNA synthesis and RT-PCR

RNA was extracted from cell bodies and endfeet using a combination of Trizol LS and RNeasy Plus Micro kit (Qiagen, 74034). Samples were thawed on ice and mixed by vortex for 15sec each. They were incubated at room temperature for 5min. 133µl chloroform (per 500µl Trizol LS) was added and samples were shaken vigorously for 15sec. Samples were incubated at room temperature for 2min and then centrifuged at 12,000g for 15min at 4°C. The aqueous phase was removed and mixed with an equal volume of 70% ethanol in DEPC water. The entire sample was transferred to the RNeasy MiniElute spin column (from Qiagen kit) and processed following the manufacturer’s instructions. RNA was eluted in 15µl RNase-free water and stored at - 80°C until use. cDNA was synthesized using an iScript kit according to instructions (Bio-Rad, 1708890). RT-PCR was performed with GoTaq Green Master Mix (Promega, M7122). Primers and annealing temperatures provided in Table S11.

### RNA sequencing and bioinformatic analysis

For each sample, brains from multiple litters were pooled. Endfoot replicates were at least 300,000 endfeet from 3-4 litters. Cell body replicates contained 50,000 cells per litter from 3-4 litters. 3 biological replicates for both endfeet and cell bodies were sequenced. cDNA libraries were produced using an ultra-low input kit (Takara, 634888). Samples were sequenced on a NovaSeq 6,000 S-Prime sequencer with 150bp paired-end reads. Libraries were sequenced to a read depth of 100-126 million reads per sample. Reads were mapped to the reference genome, GRCm38, using STAR (vSTAR.2.6.1d). Counts were quantified with Salmon (v0.14.1). Differential expression between cortex and endfoot samples was calculated using Deseq2 (v1.26.0). Mouse genome vM20 annotation. Initial Bioinformatic analysis performed by Jianhong Ou, PhD.

Comparisons of sequence length and GC content were performed using FeatureReachR as available on GitHub (https://github.com/TaliaferroLab/FeatureReachR) [41]. Case condition was endfoot enriched transcripts (FC >10, average FPKM in endfeet >10) control condition was cell body genes (log_2_(FC) < 0, average FPKM in cell body >10). Gene class assignment performed with PANTHER Classification System using Biological Processes categories (https://pantherdb.org/). Endfoot transcriptome overlap with disease, synapse, and PAP datasets were represented as area-proportional Venn diagrams created in BioVenn (https://www.biovenn.nl/index.php) [92]. Gene Ontology analysis for gene class enrichment performed using The Gene Ontology Resource and biological processes categories (https://geneontology.org/) [93, 94]. 500 endfoot enriched genes with the highest FC compared to cell body were used as input.

Bootstrap analysis was used to test the significance of overlapping transcripts observed between Endfoot localized and enriched transcripts and previously reported datasets. Endfoot localized and endfoot enriched data sets were compared to a randomized data set with an equal number of transcripts. Random lists were generated from all transcripts identified by RNA-seq in cell body and/or endfoot samples. For each comparison, 100 random lists were generated and the average overlap of the 100 random lists with the dataset of interest was reported. Statistical significance was assessed by two-tailed Fisher’s exact test.

### Immunofluorescence staining (IF)

Mouse embryonic brains were fixed overnight in 4% paraformaldehyde (PFA)/PBS, rinsed in PBS, and incubated in 30% sucrose/PBS overnight, before embedding in NEG-50 (Epredia, 6502) for cryopreservation. Cryostat coronal sections (20μm or 40μm) from the somatosensory cortex were then generated for immunofluorescence staining. Briefly, sections were permeabilized with 0.3% Triton X-100/PBS and blocked with 5% normal goat serum (NGS)/PBS at room temperature (RT). Sections were incubated with primary antibodies overnight at 4°C and species-specific secondary antibodies (Invitrogen, Alexa Fluor conjugate, 1:800) and Hoechst (Invitrogen, 33342, 1:1000) for 30min to 2h at RT. Finally, sections were mounted with Vectashield antifade mounting medium (Vector Labs, H-1000). The following primary antibodies were used: anti-Myc (Cell Signaling, 2278, 1:100), anti-HA (Roche, 11867423001, 1:500), anti-p150^Glued^ (BD Biosciences, 610473, 1:50), anti-Laminin (Millipore, AB2034, 1:500), anti-p73 (Cell Signaling, 14620, 1:250) and anti-LHX6 (Santa Cruz, sc-271433, 1:500). Images were acquired using a Zeiss Axio Observer Z1 microscope equipped with an Apotome or an Andor Dragonfly Spinning Disk Confocal microscope. Image measurements and quantifications were blindly performed using Fiji (ImageJ) or QuPath.

### Single-molecule inexpensive fluorescence *in situ* hybridization (smiFISH)

Mouse samples were cryopreserved and sectioned as described above for IF and smiFISH was performed as previously described [46]. Ferret samples were collected as floating sections, transferred to charged slides, and incubated at 37°C for 4 hours in a hybridization oven. Samples were cooled, fixed in 4% PFA in DEPC 1xPBS for 30 minutes at room temperature, and washed twice in DEPC 1xPBS. Finally, sections were treated with Proteinase K (10μg/mL New England Biolabs P8107S) in DEPC 1x PBST (0.1% Tween 20) for 5 minutes at room temperature. An equimolar mix of probes (36 probes for *Dync1h1*, 24 probes for all other genes) were hybridized to FLAP X-Cy3 (IDT). Cortical sections were permeabilized with 0.5% Triton-X-100 in DEPC PBS, rinsed with smiFISH wash buffer (10% formamide/2X SSC/DEPC water), and incubated with the hybridized probes (1:100 for *Dync1li2*, *Dynll2*, *Dync1h1*, and *Dctn1*, or 1:50 for *Gab2*, *Mmp14*, and *Sh3pxd2a*) in hybridization buffer (10% formamide/2X SSC/10% Dextran Sulfate Sodium Salt/DEPC water) overnight at 37°C. Samples were then rinsed in smiFISH wash buffer, incubated with Hoechst, rinsed in DEPC PBS, and mounted with Vectashield. Images were acquired with a Zeiss Axio Observer Z1 microscope equipped with an Apotome and smiFISH signal intensity quantifications performed in Fiji (ImageJ). Probe sequences provided in Table S11.

### siRNAs and *in utero* electroporation (IUE)

A mixture of four siRNAs targeting the coding sequence and the 3′ UTR were used to effectively depleted *Dync1li2* transcripts. siRNAs were validated in N2a cells by RT-qPCR. siRNA sequences provided in Table S11. Pregnant CD1 dams were kept under anesthesia with an isoflurane vaporizer during the entire surgical procedure. Mouse embryonic brains were injected with 1μl of solution, consisting of plasmid or siRNAs of interest and Fast Green FCF dye (Sigma, 861154) in sterile water or PBS, and electroporated using 5 pulses at 60 to 70V for 50ms at 950-ms intervals with Platinum BTX Tweezertrodes.

### Breasi-CRISPR genome-editing

Breasi-CRISPR method was used to epitope-tag endogenous proteins in the developing mouse brain. It was performed as previously described [95, 96]. The crRNAs and single-stranded oligodeoxynucleotide (ssODN) HDR templates were designed on the IDT website. Briefly, the tracrRNA (IDT, 1072532) and crRNA were hybridized (1:1) at 98°C for 2min followed by 10min incubation at RT, resulting in the final gRNA. A solution of 3μl gRNA (100μM), 4μl HDR template (100μM), 1μl Cas9 nuclease (IDT, 1081058), 1μg/μl pCAG-EGFP, 1μl Fast Green, and water to 10μl was prepared, incubated at 37°C up to 30min, and delivered to E14.5 mouse embryonic brains by IUE. tracrRNA and crRNA sequences provided in Table S11.

### RGC basal process branching complexity and endfoot organization analyses

RGC branching complexity was assessed as previously described [18, 19]. For quantification of basal process branches, RGCs were sparsely labelled with pGLAST-EGFP-CAAX introduced by IUE. Using a Zeiss 780 Inverted Confocal microscope or an Andor Dragonfly Spinning Disk Confocal microscope, 20-40μm Z-stack images were taken at 63X, 0.25μm per step. Next, 3D projections were generated for individual cells in ImageJ with the following parameters-Projection method: Brightest Point, Axis of rotation: Y-Axis, Slice spacing: 0.2, Initial angle: 0, Total rotation: 360, Rotation angle increment: 5, Lower transparency bound: 1, Upper transparency bound: 255, Opacity: 0, Surface depth-cueing: 100, Interior depth-cueing: 50, Interpolate: checked. We assigned branches to first, second, third, fourth, fifth, sixth or seventh order based on the number of ramifications between the basal process and the branch of interest. Projections >5μm were considered as branches. Round-shaped branch ends thicker than the processes were considered endfeet and categorized as attached, unattached, or protruded relative to the Laminin staining of the BM.

The RGC-branching thickness represents the measurement from the BM, labeled by Laminin, to the first ramification of the basal process (i.e., the initial branching point of an individual cell). The MZ was identified by the sparse nuclei between the pia and layer 2 of the CP for measurements of MZ thickness and number of cells within the MZ. For quantification of cells, 5μm Z-stacks images were taken at 20X, 1μm per step, using a Zeiss Axio Observer Z1 microscope equipped with an Apotome, and maximum intensity projections made in ImageJ. Positive cells were then counted using QuPath in 300μm wide ROI areas; background signal above the pia or in blood vessels counted as cells were manually excluded.

### LOCAL-KD

#### Plasmids

pCAG-NLS-Cas13d-NLS-HA-T2A-EGFP: U6-NLS-Cas13d-NLS-HA-T2A-EGFP plasmid (Addgene #109049) was used to amplify the NLS-Cas13d-NLS-HA-T2A-EGFP sequence which was inserted into an empty pCAG backbone. pCAG-Cas13d-HA-T2A-EGFP-ELS: Cas13d and HA-T2A-EGFP sequences were amplified from Addgene #109049 (removing the NLSs) and inserted into an empty pCAG backbone. Then the 5’UTR of mouse Arhgap11a was added as an Endfoot Localizing Sequence (ELS) following the EGFP. U6-gRNA: Addgene #109053. gRNAs were annealed and inserted using built in BbsI sites. pCAG-gRNA: The RfxCas13d DR30 sequence and BbsI sites were amplified from Addgene #109053 and inserted into an empty pCAG backbone. gRNAs were annealed and inserted using BbsI sites.

#### gRNA design and validation

Cas13 gRNAs were designed using the cas13design tool (https://cas13design.nygenome.org/) [97, 98]. Sequence inputs used from NCBI: RefSeq accession numbers NM_009415.3 (*Tpi1*) and NM_001013380.2 (*Dync1li2*). *Dync1li2* gRNAs were selected for those targeting the CDS as UTRs are predicted to diverge between transcript variants. gRNA sequences detailed in Table S11. Top scoring gRNAs were validated *in vitro* in N2A cells before being used *in vivo*.

Plated 75,000 N2A cells per well of a 12 well plate. Cells were incubated overnight at 37°C. Media was changed 1 hour prior to transfection. DNA was transfected with Lipofectamine 2000 (ThermoFisher 11668019) according to manufacturer’s instructions. For each well, 3µl Lipofectamine 2000, 1μg pCAG-NLS-Cas13d-NLS-HA-T2A-EGFP, 1μg U6-gRNA, and 60μl Opti-MEM low serum media (Gibco 31985070) were added. Lipofection mix was incubated on the cells for 5 hours at 37°C and then replaced with fresh DMEM (ThermoFisher 11965092). Cells were collected in 500μl Trizol (ThermoFisher 15596026) 24 hours post transfection.

### *In vivo* LOCAL-KD and validation

1-1.5μg pCAG-NLS-Cas13d-NLS-HA-T2A-EGFP or pCAG-Cas13d-HA-T2A-ELS were introduced by IUE with 1μg U6-gRNA (NLS) or pCAG-gRNA (ELS). Control condition was Cas13d alone with no gRNA. For sorting and qPCR experiments, 0.5μg pCAG-EGFP was added. For morphology analysis, 0.5μg pCALNL-mScarlet-CAAX and 1ng pCAG-Cre were added. IUE was performed as previously described in CD1 mice using 60V electroporation at E15.5. Samples were collected 24 hours later at E16.5.

For qPCR validation of LOCAL-KD, endfeet and cell bodies were sorted as described above using the EGFP introduced by IUE, instead of *Nestin*-EGFP, to mark EGFP positive endfeet and cell bodies. qPCR was performed using iTaq SYBR green master mix (Bio-Rad 1725124) and transcript specific primers detailed in Table S11. Reactions run on a QuantStudio 6 Real-Time PCR system (ThermoFisher). For *Tpi1* qPCR annealing temperature 58°C and normalized to *Ddr2* levels for endfeet and β-actin for cell bodies. For *Dync1li2* qPCR annealing temperature 60°C and normalized to *Ccnd2* levels for endfeet and β-actin for cell bodies.

Samples were cryopreserved and sectioned as described above for IF and smiFISH. Sections were thawed at RT for 20 minutes, washed in DEPC 1xPBS, permeabilized with 0.5% triton in DEPC PBS for 30 minutes, and washed again in DEPC PBS. RNse free BSA solution was prepared by dissolving nuclease free BSA (VWR 97061-420) in DEPC 1xPBS, filtering with a 0.22µm syringe filter, and treating with RNse OUT (ThermoFisher 10777019) 1:250 for 10 min at RT. Incubated with anti-HA primary antibody (Roche) in 1% BSA at RT for 1 hour, washed DEPC 1xPBS, incubated with secondary antibody (anti-Rat Alexa Fluor 350, Invitrogen A-21093) in 1% BSA at RT for 30 minutes. Performed smiFISH protocol as described previously beginning with the washes in FISH wash buffer. Mouse *Dync1li2* smiFISH probes used 1:100. Imaged at 40x, 10µm Z-stack with 1µm step size on a Zeiss Axio Observer Z1 microscope equipped with Apotome. Maximum intensity projections made in ImageJ.

#### smiFISH-IF quantification

Quantification was performed on maximum intensity projections in ImageJ. EGFP signal was used to draw ROIs around all individual endfeet. Measured the total area and % *Dync1li2* smiFISH positive area for each ROI (endfoot). Calculated the *Dync1li2* smiFISH positive area for each ROI. Each ROI was classified as Cas13d positive or negative based on HA signal from the smiFISH-IF. 2-4 sections quantified per brain. For each section, the % *Dync1li2* positive area in the Cas13d positive ROIs was normalized to the Cas13d negative ROIs. Plots represent average of all sections per brain normalized to the control condition.

#### Sparse labeling

For sparse labeling of RGCs with LOCAL-KD (Fig. 7) 0.5 µg/µl pCALNL-mScarlet-CAAX and 1 ng/ µl pCAG-Cre were added to the Cas13d and gRNA described above. Embryos were IUE’d at E15.5 and samples were collected 24 hours later at E16.5.

## Acknowledgements

We thank the members of the Silver Lab for helpful discussions and careful reading of the manuscript. We also thank the Duke FACS, Sequencing, and Light microscopy Core Facilities. We thank Qiang Lu for sharing mice, Jianhong Ou for RNA-seq analysis and RJ Madsen for bioinformatic assistance. We thank LJ Pilaz for initial investigations of endfeet and advice on BREASI-CRISPR.

## Author contributions

Conceptualization: BRD, CMM, and DLS. BRD developed the endfoot purification method, performed RNAseq experiments, FeaturereachR data analyses, disease genes, venns, initial validation of targets and developed the LOCAL-KD method. CMM performed Dynein component validation and *Dync1li2* siRNA KD functional studies. CFL performed 3D reconstruction imaging, *Tpi1* smiFISH, and *Cas13d* smiFISH. LDS performed RNA stability and ribosomal component analysis. SPS assisted with optimization of LOCAL-KD. VF and VB provided ferret tissue. Writing original draft: BRD, CMM, and DLS. All authors edited the manuscript. Funding acquisition and project oversight: DLS.

---

**Supplemental Fig 1.**
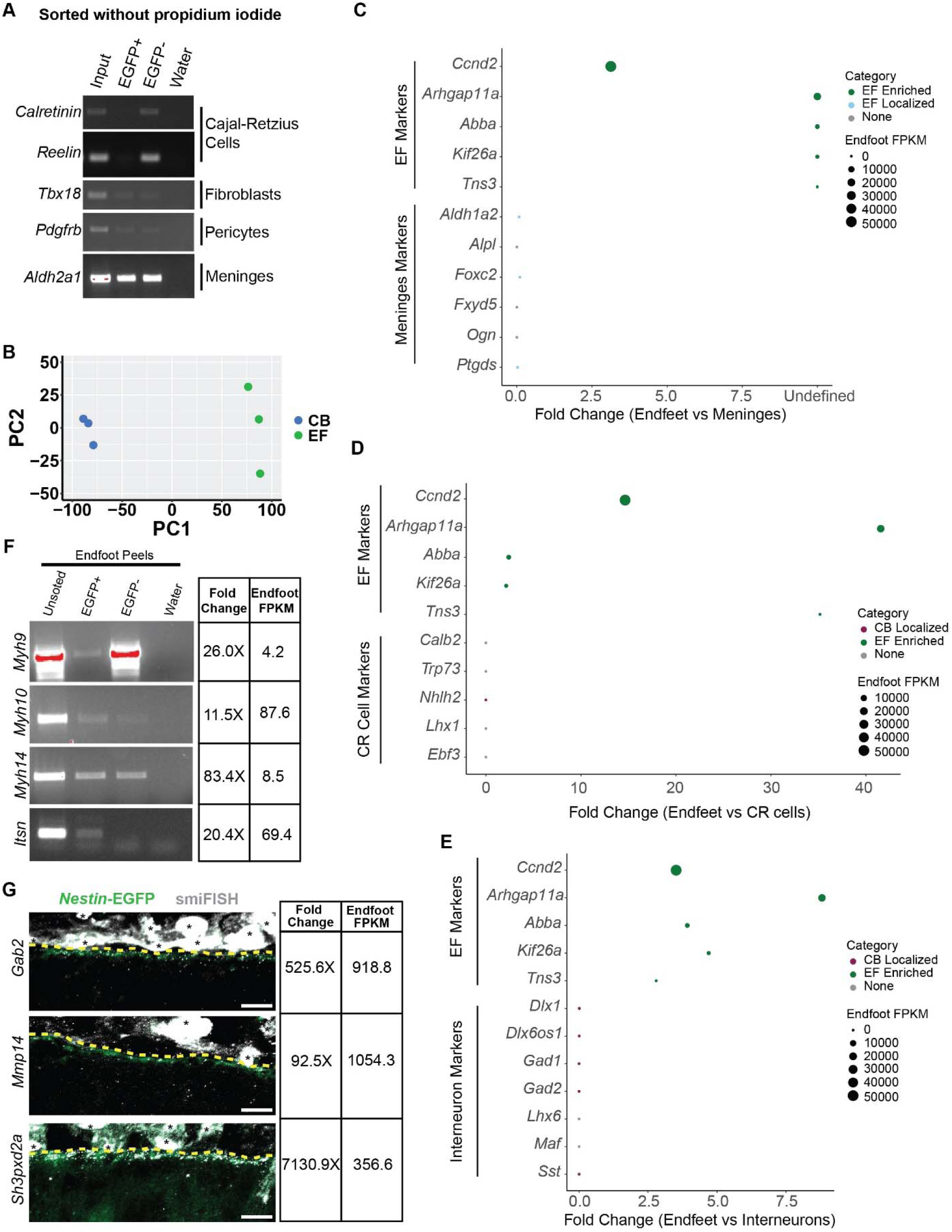
qPCR and FISH validation for purification of RGC subcellular structures from embryonic cortices. **(A)** RT-PCR for marker genes (Left) of cell types (Right) potentially contaminating sorted endfeet. Comparison of expression in the input, EGFP+, and EGFP-samples for endfeet sorted without propidium iodide. **(B)** PCA plot comparing biological replicates for the cell body (CB, blue) and endfoot (EF, green) RNA sequencing samples. **(C-E)** Comparison of known endfoot markers and markers of meninges (C) [23], CR Cells (D) [24], or interneurons (E) [25] in sorted endfeet. Circle size represents average FPKM of each transcript in endfeet. Circle color denotes classification category assigned to each transcript (Endfoot Enriched-green, Endfoot Localized-light blue, Cell Body Localized-maroon, or None-grey) based on FPKM in endfeet and enrichment in endfeet vs cell bodies. Endfoot markers that were not detected in meninges have a reported fold change of “undefined” as the endfoot FPKM is divided by zero (C). **(F)** RT-PCR validation of select endfoot localized transcripts in purified endfeet (left) of varying fold change and average FPKM in endfeet (right). **(G)** smiFISH (grey) validation of select endfoot localized transcripts localization to endfeet (green) (left) of varying fold change and average FPKM in endfeet (right). Yellow dotted line denotes boundary between endfeet and the pia. Black asterisks denote autofluorescence background from the meninges. Scale bars: 10µm.

**Supplemental Fig 2.**
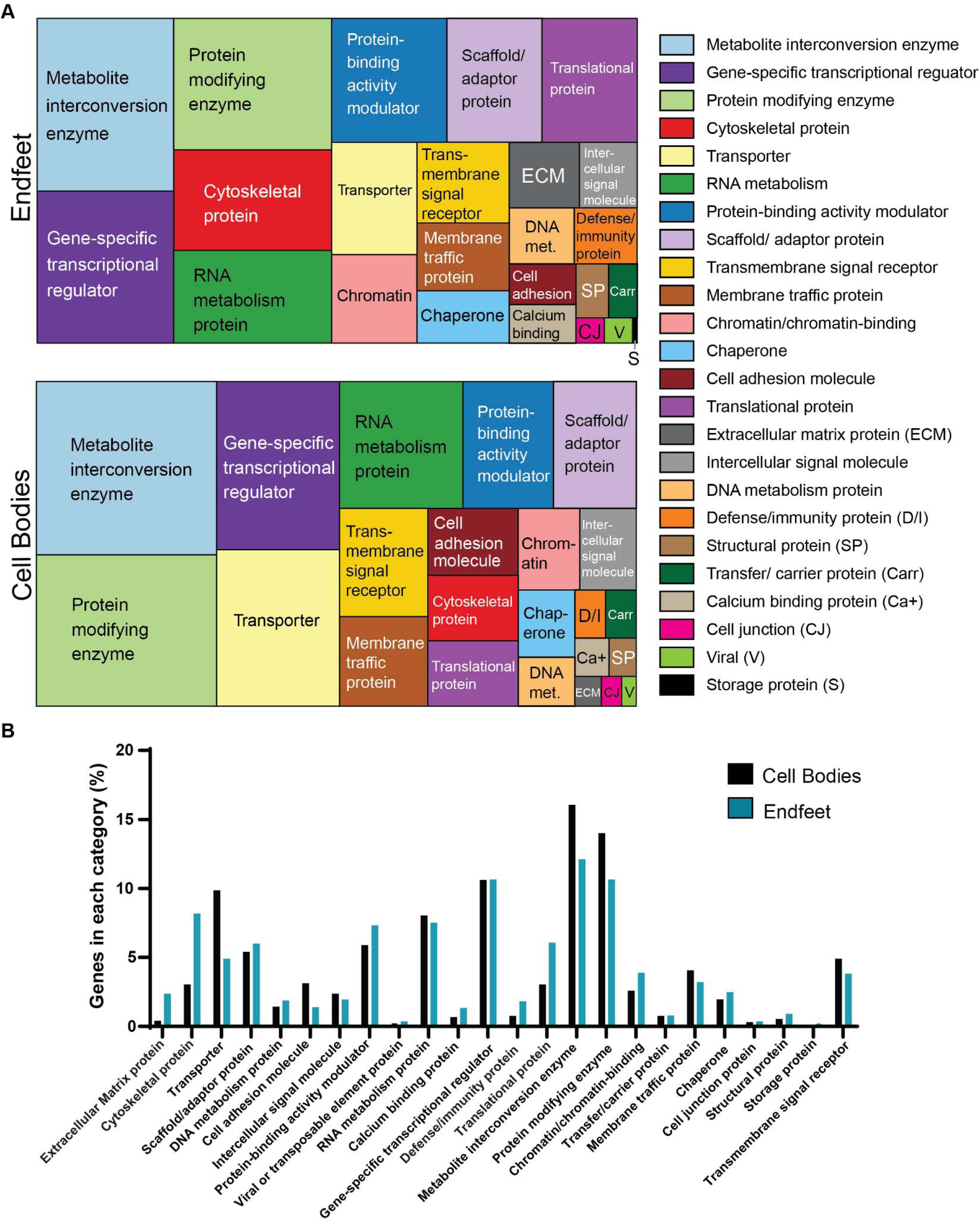
PANTHER analysis of endfoot and cell body localized transcripts. **(A)** Treemaps of GO analysis for protein classes represented in the cohort of endfoot localized (top left) and cell body localized (bottom left) genes. Key and full names of protein class categories represented in treemaps (right). **(B)** Bar graph comparing proportion of genes in each class between the endfoot localized (teal) and cell body localized (black) transcriptomes.

**Supplemental Fig 3.**
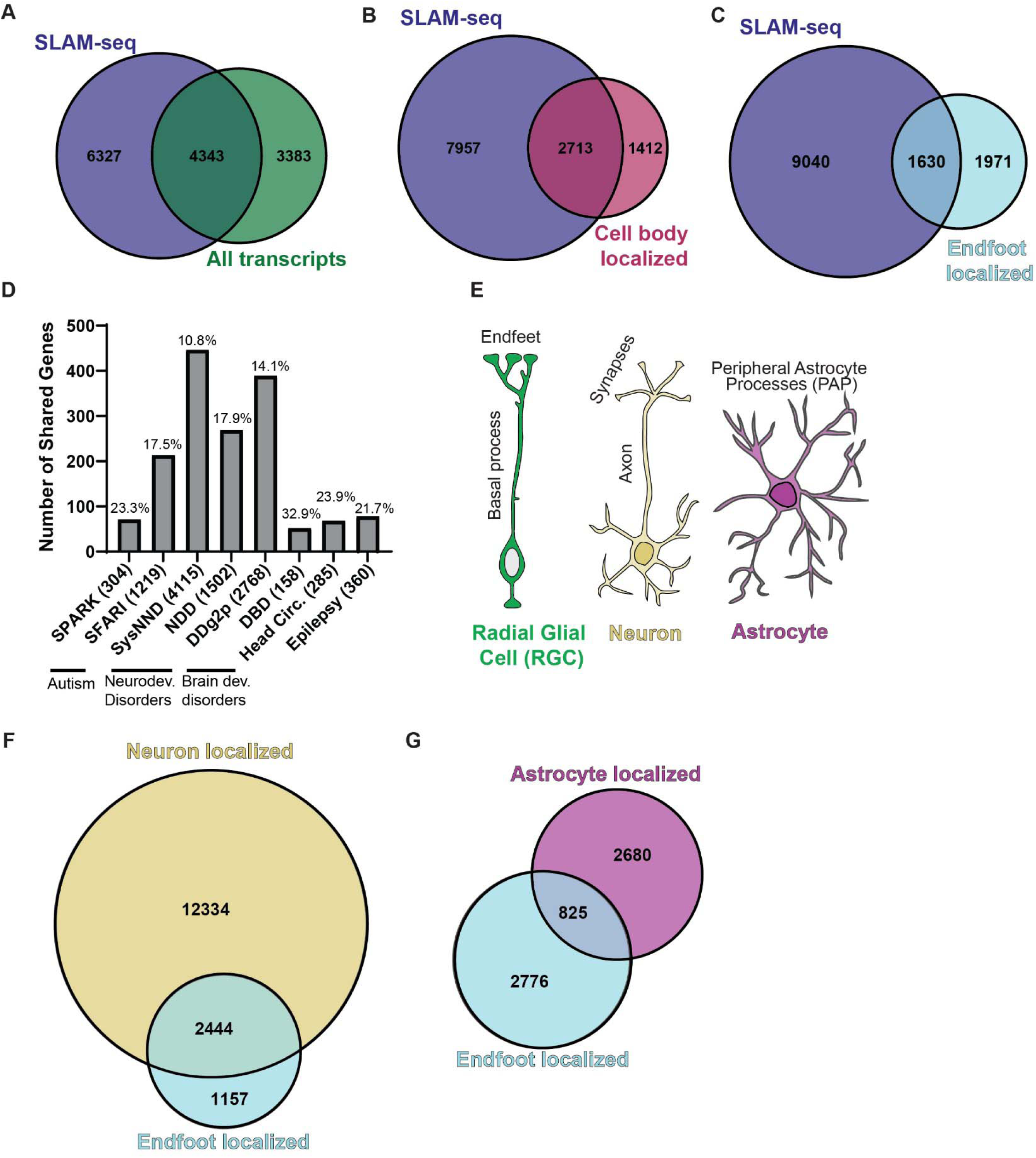
Analysis of mRNA stability and overlap of endfoot localized transcripts with neurons and astrocytes. **(A)** Comparison of transcripts represented in SLAM-seq data (blue) and the cell body and endfoot samples (green). **(B)** Comparison of transcripts in SLAM-seq (blue) and cell body localized (maroon). **(C)** Comparison of transcripts in SLAM-seq and endfoot localized datasets (light blue). **(D)** Genes in neurodevelopmental disease databases that are also localized to RGC endfeet. Parentheses denote total number of genes reported in each disease database. Numbers above bars report % of database genes that overlap with the endfoot localized transcriptome. **(E)** Cartoon representation of the complex morphology of radial glial cells (green), neurons (yellow), and astrocytes (magenta) with their subcellular structures of interest (endfeet, synapses, and peripheral astrocyte processes) labeled. **(F)** Venn diagram comparing transcripts identified as endfoot localized (light blue) to previously published data sets of transcripts localized to neuronal synapses (yellow). **(G)** Venn diagram comparing transcripts identified as endfoot localized (light blue) to previously published data sets of transcripts localized to peripheral astrocyte processes (PAPs) in astrocytes (magenta).

**Supplemental Fig 4.**
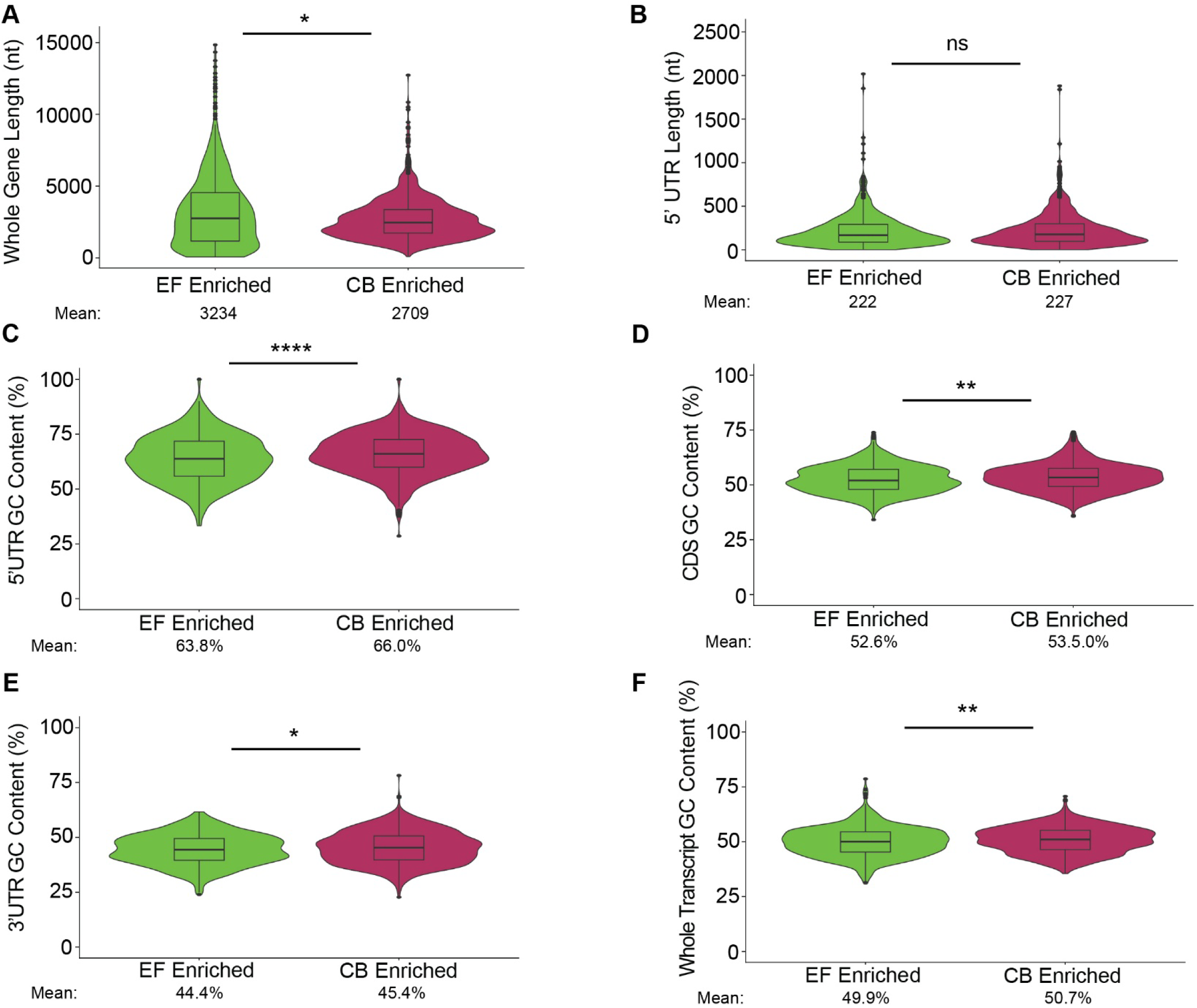
Bioinformatic analyses of transcriptomic datasets. **(A, B)** Violin plots comparing the length of the whole transcripts **(A)** and 5′ UTRs **(B)** between endfoot enriched (EF enriched) transcripts (green) and cell body enriched (maroon). **(C-F)** Violin plots comparing the GC content of the 5′ UTRs **(C)**, coding sequence (CDS) **(D)**, 3′ UTRs **(E)** and whole genes **(F)** between EF enriched transcripts (green) and cell body enriched (maroon). Mean for each condition is noted under the graph (A-F). Statistics: (A-F) Wilcoxon rank-sum test. ns p-value > 0.05, * p-value < 0.05, **p-value < 0.01, *** p-value < 0.001, **** p-value < 0.0001.

**Supplemental Fig 5.**
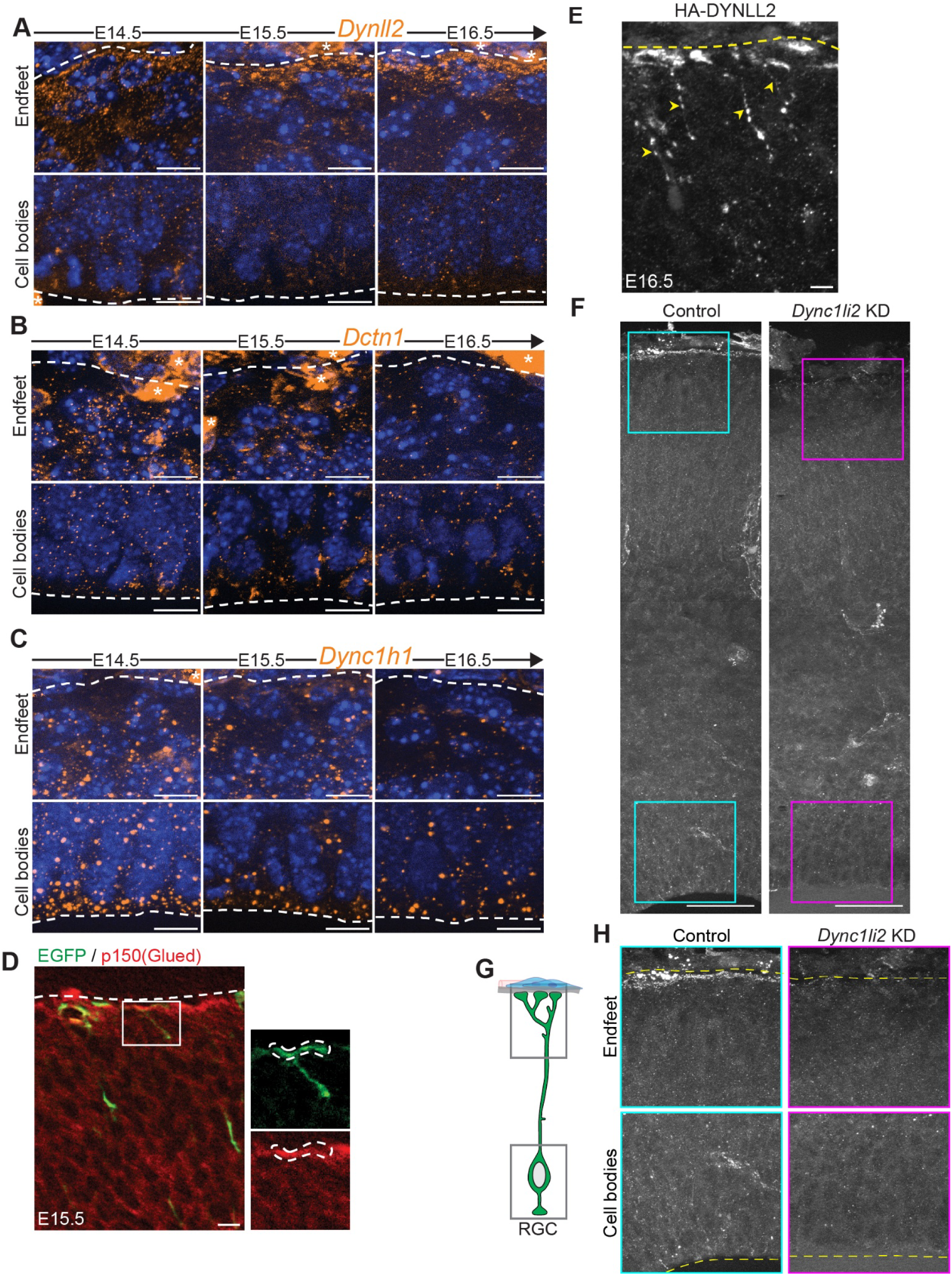
Dynein-dynactin subunits *Dynll2*, *Dctn1*, and *Dync1h1* localization to RGCs during cortical development. **(A-C)** Localization of **(A)** *Dynll2* **(B)** *Dctn1* and **(C)** *Dync1h1* smiFISH puncta (orange) to RGC endfeet (top) and cell bodies (bottom) from E14.5 to E16.5 in the embryonic mouse cortex. Nuclei labeled with DAPI (blue). White dotted lines indicate pial (top) and ventricular (bottom) borders and asterisks denote background signal from the meninges or blood vessels**. (D)** Immunofluorescence depicting p150Glued (red) in EGFP-electroporated RGCs. Right panels highlight colocalization in RGC basal endfoot (dotted lines) at E15.5. **(E)** Localization of tagged Myc-DYNLL2 in RGC endfeet and basal processes at E16.5. Yellow dotted lines denote pial boundary. Yellow arrow heads mark tagged protein in basal processes. **(F)** Representative images showing *Dync1li2* smiFISH signal in the entire cortex in control and *Dync1li2* KD brains. Boxes outline ROIs magnified in (H). **(G)** RGC cartoon with areas magnified in (H) outlined (grey). **(H)** Magnified images of the pial and ventricular areas, highlighting striking *Dync1li2* KD in the RGC endfeet and discrete KD in the RG cell bodies compared to control (left, blue). Scale bars: (A-E) 10µm, (F) 50µm. smiFISH, single-molecule inexpensive fluorescent in situ hybridization; RGC, radial glial cell; ROI, region of interest.

**Supplemental Fig 6.**
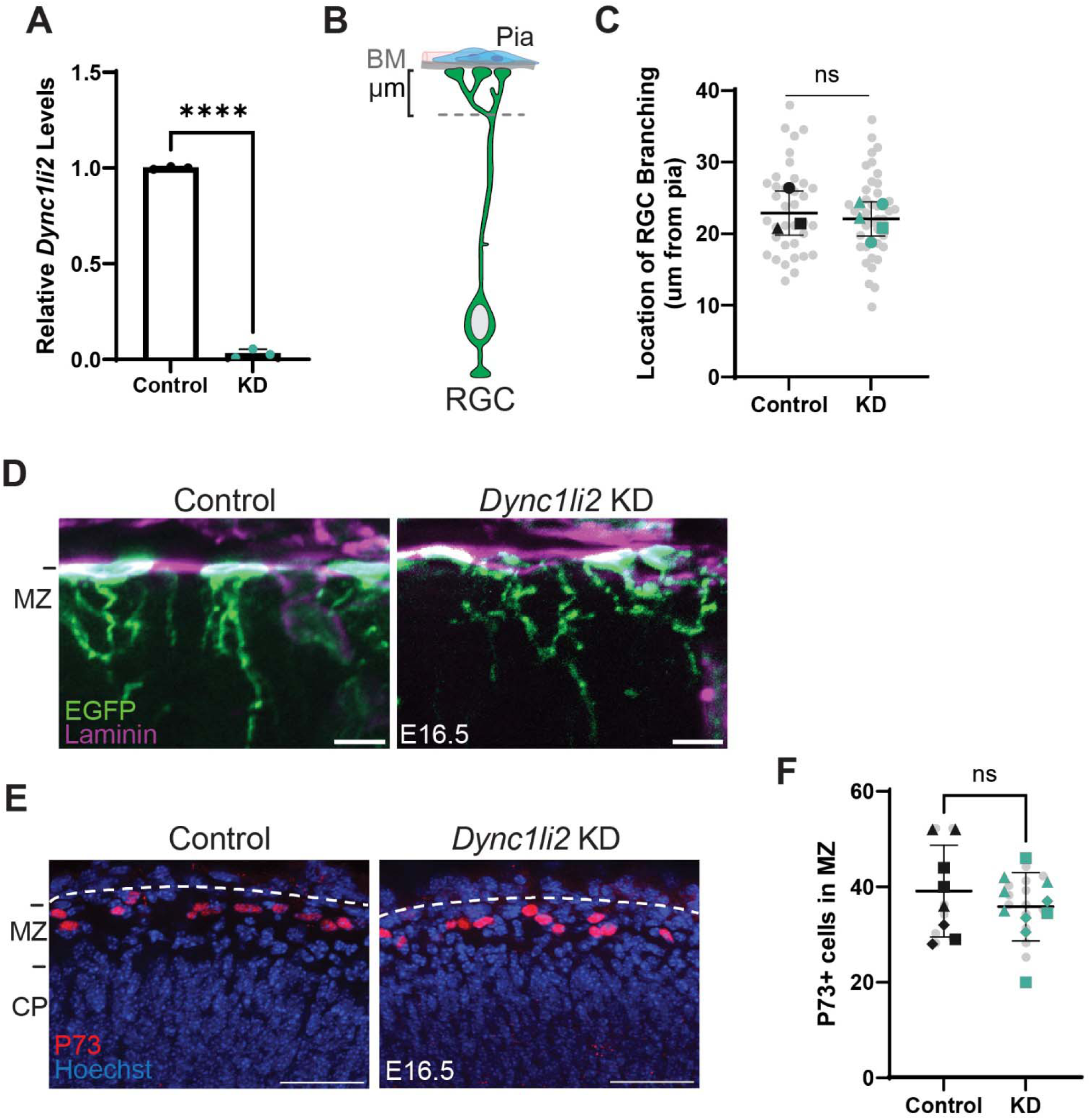
Analysis of RGC morphology and Cajal-Retzius cells in *Dync1li2* KD brains. **(A)** qPCR quantification of Dync1li2 levels in sorted endfeet 24 hours after IUE with scrambled (Control) or *Dync1li2* (KD) siRNA. n= 3 biological replicates. Each replicate contains 20,000-30,000 endfeet from 3-5 brains. **(B)** Cartoon representation of RGC-branching thickness measuring criteria. **(C)** Graph depicting branching thickness quantifications. n=36 cells, 3 control brains, and n=42 cells, 5 KD brains, from 3 litters **(D)** Representative images of EGFP-labeled RGC endfeet (green) attachment to the BM stained with Laminin (magenta) at E16.5. **(E)** Representative images of P73 staining (red) to label CR cells and Hoechst (blue) at E16.5. White dotted lines mark pial boundary. MZ and CP (left) denote MZ region quantified in (F). **(F)** Quantification of P73+ cells in the MZ in control (black) and *Dync1li2* KD (teal) cortical sections. n= 9 sections, 8 control brains, and n= 15 sections, 10 brains, from 3 litters. Scale bars: (D) 10 µm, (E) 50µm. Super-plots depict ROIs in gray dots and individual data representing different brains in black (control) or teal (KD) dots; litters are coded by shape. (A, C, F) Student unpaired, two-tailed t test; Error bars: SD ns p-value > 0.05 BM, basement membrane; CR cells, Cajal-Retzius cells; KD, knockdown; MZ, marginal zone.

**Supplemental Fig 7.**
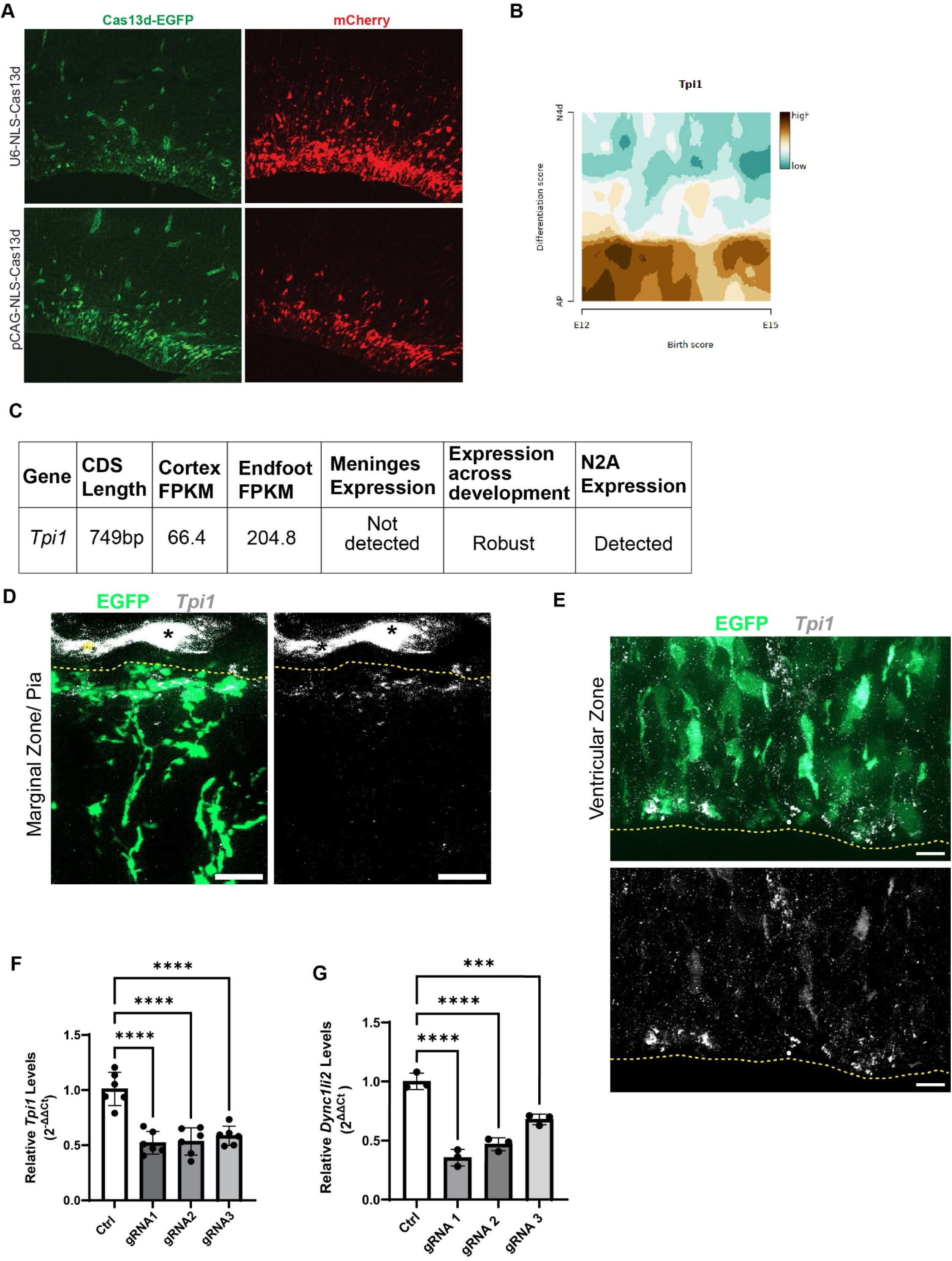
Analysis of LOCAL-KD efficacy *in vitro* and *in vivo*. **(A)** Comparison of NLS-Cas13-NLS-HA-T2A-EGFP expression (left) when driven by U6 (top) or CAG (bottom) promotors compared to electroporation efficiency (red mCherry right). **(B)** Expression of *Tpi1* in compared to cell differentiation score (AP-N4d) and birth score (E12-E15) [99]. **(C)** Table of characteristics considered when selecting *Tpi1* as a proof-of-concept gene. **(D-E)** SmiFISH (grey) for *Tpi1* in endfeet (D) and cell bodies (E). RGC morphology marked by electroporated EGFP (green). Yellow dotted lines denote pial (D) and ventricular (E) boundaries. Asterisks mark background signal from blood vessels **(F-G)** *Tpi1* (F) and *Dync1li2* (G) levels in N2A cells transfected with NLS-Cas13d and different gRNAs for validation *in vitro*. (F-G) Points represent individual wells of N2A cells. Statistics: One-way ANOVA ***=p<0.0001, ****=p<0.00001 Error Bars: SD.

**Supplementary Table 1** Subcellular transcriptome of RGCs

**Supplementary Table 2** Panther analysis of endfoot localized transcripts

**Supplementary Table 3** Half-lives of subcellularly localized transcripts

**Supplementary Table 4** Overlap of endfoot localized transcripts with known disease loci

**Supplementary Table 5** Overlap of endfoot localized transcripts with known localized endfoot RNAs and proteins

**Supplementary Table 6** Overlap of endfoot localized transcripts with known localized transcripts of neurons and astrocytes

**Supplementary Table 7** Overlap of endfoot localized and enriched transcripts with ribosomal genes

**Supplementary Table 8** Overlap of endfoot enriched transcripts with known disease loci

**Supplementary Table 9** Overlap of endfoot enriched transcripts with known localized transcripts of neurons and astrocytes

**Supplementary Table 10** Gene ontology of endfoot enriched transcripts

**Supplementary Table 11** Reagents used in this study

